# The medial prefrontal cortex encodes procedural rules as sequential neuronal activity dynamics

**DOI:** 10.1101/2025.01.16.633469

**Authors:** Shuntaro Ohno, Masanori Nomoto, Kaoru Inokuchi

## Abstract

The prefrontal cortex plays a crucial role in procedural rule learning; however, the specific neuronal mechanism through which it represents rules is unknown. We hypothesized that sequential neuronal activities in the prefrontal cortex encode these rules. To investigate this, we recorded neuronal activities in the medial prefrontal cortex of mice during rule learning using Ca^2+^ imaging. We utilized a method based on convolutional negative matrix factorization, iSeq, to automatically detect temporal neuronal sequences in the recorded data. As rule learning advanced, these neuronal sequences began to encode critical information for rule execution. In mice that had mastered the rule, the dynamics of neuronal sequences could predict success and failure of reward acquisition. Furthermore, the composition of cell populations within the neuronal sequences was rearranged throughout the learning process. These findings suggest that as animals learn a rule, the medial prefrontal cortex continually updates its neuronal sequences to assign significance to behavioral actions crucial for reward acquisition.

## Introduction

For animals to survive, it is crucial that they understand their current situation, take appropriate actions, and secure rewards such as food and safety. To efficiently obtain rewards, it is necessary to extract important pieces of information from external stimuli received in the past and actions previously taken, and to integrate these into an appropriate sequence that leads to reward acquisition. This order constitutes a procedural rule, and the individual stimuli and actions that make up this procedural rule are defined as elements of the rule.

The prefrontal cortex (PFC) is considered an important brain region for rule learning^1-3^. To obtain and retain procedural rule information, the PFC must be capable of representing elements of the rule. Indeed, neuronal activity in the PFC encodes various pieces of information that form elements of rules. For example, neuronal representation in the PFC can be classified into distinct clusters, each encoding specific behavior in mice^4^ and individual location-rank information for sequential working memory in monkeys^5^. In addition, reward-responsive neurons have been found in the PFC of rats^6^. It is believed that, in the PFC, important sets of information for reward acquisition are selected from these representations to form procedural rules. However, the mechanisms at the neuronal level that accomplish this process are not yet fully understood.

To explore the neural mechanisms underlying rule representation, we focused on neuronal activity patterns that appear with a consistent temporal order, known as neuronal sequences. Neuronal sequences have been identified in various brain regions across species, including the hippocampus of rats^7-9^, visual cortex of cats^10^ and mice^11^, frontal cortex of monkeys^12,13^, parietal cortex of monkeys^14^ and mice^15^, gustatory cortex of rats^16^, and premotor cortical nucleus HVC of songbirds^17^. These activities are known to correlate with behaviors that involve sequences and spans of time, making them highly compatible with the procedural aspects of rules. Furthermore, the medial PFC (mPFC) of rats contains neuronal sequences that encode movement trajectories, which reactivate upon changes in rules^18^. Therefore, we hypothesized that in the PFC, either a single neuronal sequence or the dynamics of multiple neuronal sequences represent the procedural rules.

To validate this hypothesis, we recorded neuronal activity in the mPFC of mice during procedural rule learning using Ca^2+^ imaging. We then employed the iSeq algorithm, an enhanced version of existing methods^17,19,20^, to automatically identify neuronal sequences in the recorded data. This revealed that the mPFC contained neuronal sequences that were activated in response to specific behaviors. As the mice learned the rule, the behavioral actions encoded by these neuronal sequences became more defined, allowing the dynamics of sequences to predict successful and unsuccessful reward acquisitions. Furthermore, in mice that mastered the rule, the neuronal sequences encoded information of crucial external stimuli and behavioral actions for rule execution. The fact that the cell populations comprising these neuronal sequences changed daily suggests that as animals learn rules, the PFC continuously updates its neuronal sequences to effectively encode behaviors crucial for reward acquisition.

## Results

### “iSeq” automatically detects neuronal sequences from Ca^2+^ imaging data

We attempted to measure neuronal activity in mice using Ca^2+^ imaging, aiming to automatically identify neuronal sequences in the data. To detect these neuronal sequences in substantial datasets of recorded neuronal activity, earlier studies often employed methods such as averaging neuronal activity across trials^7,8,15,18^ or using hidden Markov models^13,16^. Here, we implemented convolutive non-negative matrix factorization (ConvNMF)^19,20^, a technique that does not refer to animal behavior and facilitates the delineation of neuronal sequence patterns.

Basic negative matrix factorization (NMF)^21,22^ identifies patterns of synchronous neuronal activity^23^. This method decomposes a large two-dimensional (2D) matrix, denoted as V, which records the activity signals from hundreds of neurons across thousands of time frames, into two distinct 2D matrices: the pattern matrix W and the intensity matrix H (Figure 1A). In W, each column vector represents a synchronously activated neuronal ensemble (population activity pattern). Conversely, the rows of H indicate the temporal variation of the activation intensity for each pattern. By expanding the dimensions of W to include a time window, ConvNMF enables the representation of sequential rather than merely synchronous activity^19,20^. This method decomposes the original 2D matrix V into a 3D pattern matrix W and a 2D intensity matrix H (Figure 1B). V is then approximated by matrix U, which is reconstructed through the tensor convolution^17^ of W and H, detailed in Math Note S1. In this model, V is structured by neuron and time dimensions, W by neurons, sequence count, and sequence time window, and H by sequence count and time. By defining N, T, K, and L as the dimensions of neurons, time, sequence count, and sequence time windows, respectively, V is represented as an N × T matrix, W as an N × K × L matrix, and H as a K × T matrix. The structure of each neuronal sequence is characterized by the two dimensions in W: neurons (N) and the time window of the sequence (L). Conversely, its activity intensity over time is captured in a one-dimensional vector (row) of length T in H. Typically, ConvNMF utilizes the Euclidean distance

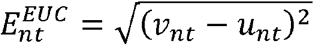

to assess the discrepancy between matrices V and U, serving as the error function. However, we opted for the Itakura-Saito (IS) divergence^24^

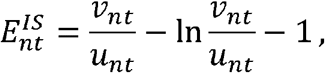

where ν _*nt*_ and *u*_*nt*_ are the activity value at time *t* of the *n*^th^ neuron in V and U, respectively (Figure 1C). We chose IS divergence because of its scale-invariant property^24^, which is particularly advantageous for analyzing data with signal-dependent noise^25^. Our recorded Ca^2+^ imaging data exhibited pronounced correlations between local mean and variance (Figure S1), indicating that the data was predominantly influenced by signal-dependent noise.

**Figure 1.**
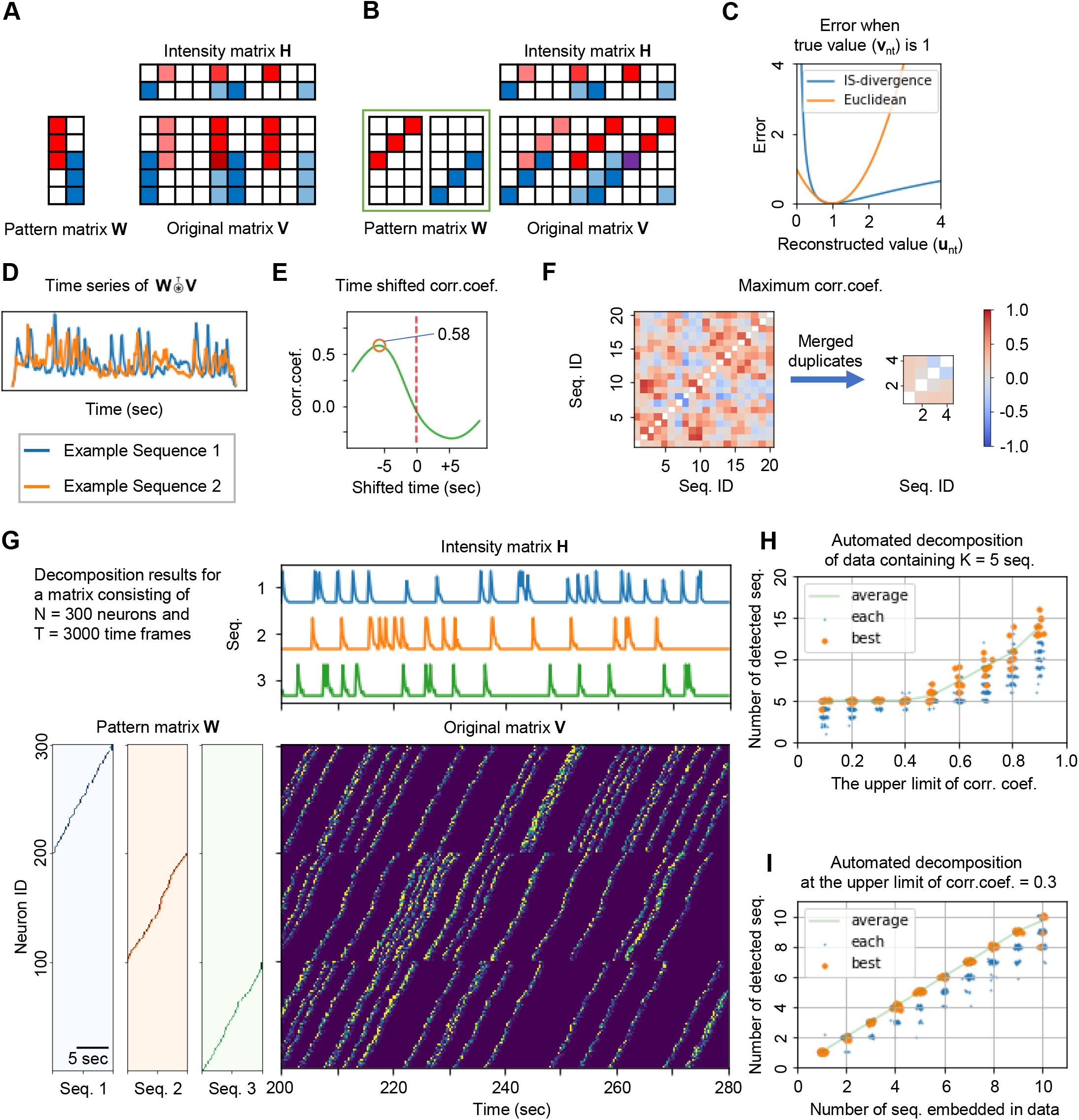
“iSeq” automatically detects neuronal sequences from Ca^2+^ imaging data. (A) Schematic for detecting synchronous activity using negative matrix factorization (NMF). (B) Schematic for detecting sequential activity using convolutional NMF. (C) Comparison of Itakura-Saito (IS) divergence and Euclidean distance. (D) Example of overlap between two neuronal sequences and the original data. (E) Example of time-shifted correlations within overlaps. (F) Similarity across all neuronal sequences (left) and similarity after merging sequences that exceed a similarity threshold (right). (G) Example of decomposing synthetic data with iSeq. (H) Relationship between the threshold setting of iSeq and the number of detected sequences in synthetic data containing K=5 neuronal sequences. (I) Correlation between the number of neuronal sequences embedded in synthetic data and the number detected by iSeq with a threshold of 0.3. See also Figures S1–S3 and Math Notes S1 and S2. Abbreviations: Corr. Coef., Correlation coefficient; Seq., Neuronal sequence.

Another important consideration when using ConvNMF is that we need to preset the number of neuronal sequences to be detected (K). If the value of K is too low, some neuronal sequences may go undetected. Conversely, if the value of K is too high, it can lead to splitting what should ideally be a single neuronal sequence into multiple parts, resulting in redundancy. Prior studies have attempted to solve this by incorporating a penalty term into the error function^17,26^. This penalty increases as the orthogonality among the detected neuronal sequences decreases, with an appropriate coefficient applied for balance. However, this method merely shifts the problem from determining K to finding the correct coefficient for the penalty term, thus making the problem more non-intuitive. Therefore, we have opted for a more direct approach.

Initially, a sufficient number of neuronal sequences were extracted. Subsequently, those determined to be identical were merged. Identity in this context is defined by the overlap (correlation) between each neuronal sequence and the original matrix V. This overlap is calculated through the transpose tensor convolution of the pattern matrix W and V (Figure 1D and Math Note S1). If the maximum value of the time-shifted Pearson’s correlation of these overlaps, calculated for two neuronal sequences, and exceeds a specified threshold (Figure 1E), these sequences are considered identical and are merged (Figure 1F). In experiments using synthetic data, the detection of an appropriate number of neuronal sequences was achieved when the threshold was set at approximately 0.3. Consequently, we consistently used a threshold of 0.3 in all subsequent calculations (Figures 1G–1I and S2A–S2B).

In addition to these methods, we developed an algorithm to ensure the convergence of the multiplicative update rules of ConvNMF (Math Note S2). We also extended the time dimension of the intensity matrix H from T to T+L−1 to accurately capture the influence of neuronal sequence activity prior to the recorded timestamps in Ca^2+^ imaging data (Figures S3A–S3B). Additionally, we compressed the original matrix V prior to decomposition to minimize computational complexity (Figures S3C–S3I). We compiled all these features into a software package designed for the automatic detection of neuronal sequences, which we named iSeq, in reference to its error function.

### The mice successfully learned the procedural rules of the Y-maze task

To investigate how the mPFC represents neuronal activity during procedural rule learning, mice were subjected to the Y-maze task (Video S1; see also STAR Methods). Each mouse was individually placed in the Y-maze, in which it could freely explore (Figure 2A, left). At the end of one arm of the maze’s branches lay the reward waiting zone, hereafter referred to as “the Zone”. Upon entering the Zone, LEDs on the maze wall were triggered to flash. If the mouse remained in the Zone for several seconds, the LEDs would switch to a continuous light, signaling the start of a 10 s period. During this time, the mouse could obtain a reward by licking a water port, referred to as “the Port”, located at the end of the trunk arm (Figure 2A, right). The mice were not explicitly taught this rule; instead, they learned through trial and error, continuing until they either acquired 40 rewards or 1 h had elapsed. The experiment spanned 6 days, with the Zone’s location switched to the opposite arm on day 4 (Figure 2B) to emphasize that the rule was procedural rather than location-specific.

**Figure 2.**
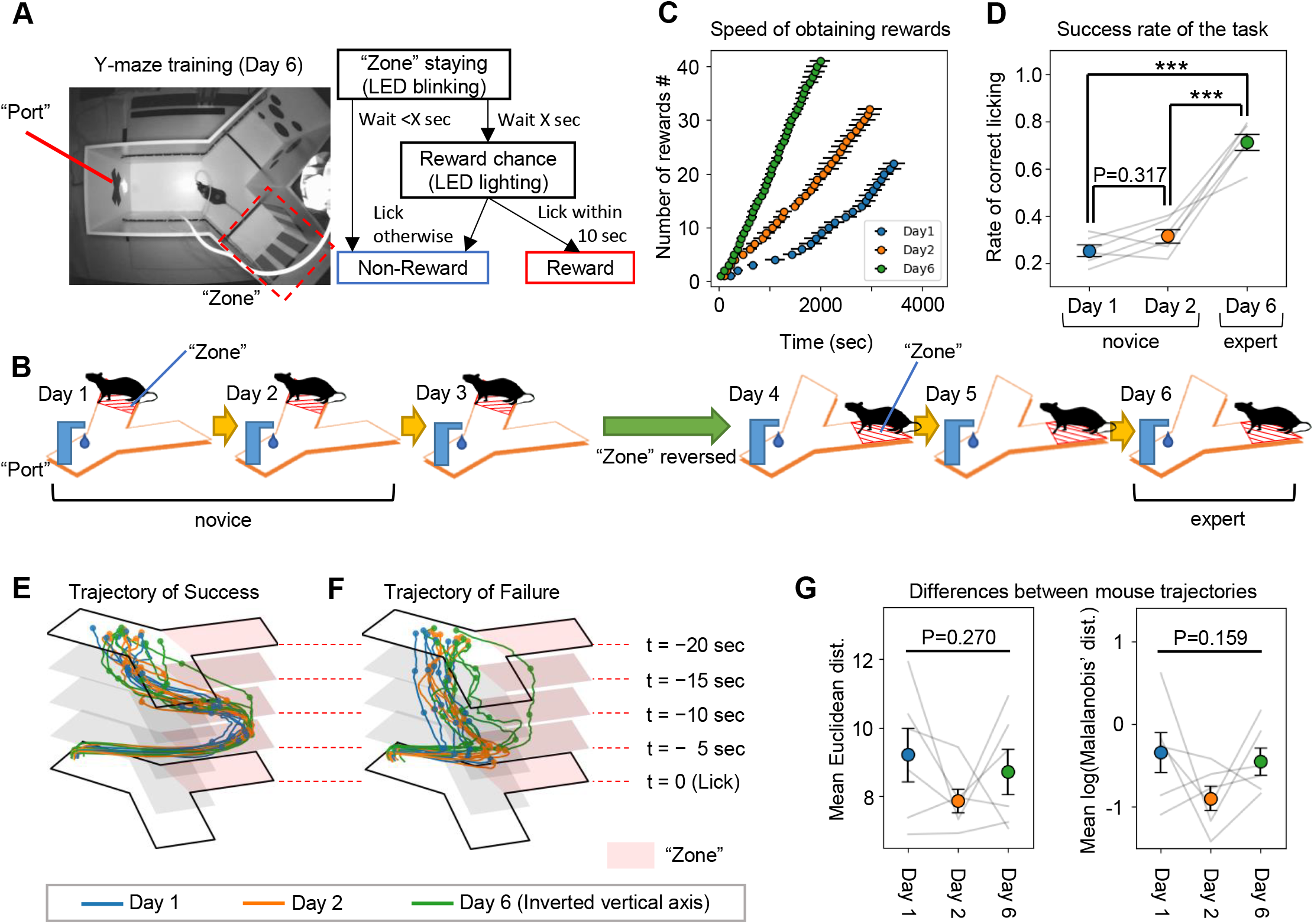
The mice successfully learned the rules of the Y-maze task. (A) Setup of the Y-maze on day 6 (left) and flowchart outlining the Y-maze task procedures (right). (B) Training schedule for the Y-maze task. (C) Pace of reward acquisition for mice on days 1, 2, and 6. (D) Success rates of reward acquisition for mice on days 1, 2, and 6 (\*\*\**P*<0.001; one-way RM ANOVA with Tukey post hoc test). (E–F) Average 20 s trajectories leading to successful (Success; E) and unsuccessful (Failure; F) reward acquisition. (G) Comparison of mouse trajectories leading to Success and Failure. The Euclidean distance of average trajectories (left) and the Mahalanobis’ distance (right), taking into account the covariance of Failure trajectories (one-way RM ANOVA). See also Figure S4 and Video S1. Abbreviations: Dist., Distance. Data are represented as mean ± SEM.

As learning progressed, the mice gradually became faster at acquiring rewards (Figure 2C), and their likelihood of obtaining a reward when licking the port also increased significantly (Figure 2D). On day 4, when the Zone’s location was reversed, there was a temporary decline in the pace of reward acquisition, yet it remained faster than on day 1 and comparable to day 2 (Figure S4A). This indicates that the mice did not perceive the task with the reversed Zone as a new challenge, but rather as a continuation of the task from the first 3 days. Furthermore, the success rate of reward acquisition on day 4 was similar to that on day 3, and there was a significant increase in the success rate from day 4 to day 5 (Figure S4B). This suggests that changing the task location on day 4 may have enhanced the mice’s understanding of the procedural rule of the Y-maze task.

Next, we examined the behavior of mice as they approached the Port during instances when they successfully licked the Port and obtained a reward (Success) and during instances when they licked the Port but failed to obtain a reward (Failure). The trajectories of the mice during Success were nearly identical on all days (Figure 2E). Additionally, the trajectories during Failure, across all days, differed significantly from those during Success (Figure 2F). When comparing these trajectory differences using either Euclidean distance or Mahalanobis’ distance (which accounts for data variance) between the average trajectories during Success and Failure, no significant differences were detected across days (Figure 2G). These results clearly demonstrate that as learning progressed, the likelihood of obtaining rewards increased, even though the trajectory of behaviors leading to successful rule completion and reward acquisition remained unchanged. Moreover, the disparity in trajectories between Success and Failure was consistent across all days.

### The mPFC contains neuronal sequences that reflect reward acquisition

To assess neuronal activity in the mPFC during procedural rule learning, we employed an in vivo Ca^2+^ imaging method to monitor 347–729 neurons from each of six mice (Figures 3A–3B). To minimize brain damage due to phototoxicity during imaging, recording sessions were limited to days 1 and 2, when the mice were ‘novices’ to the task, and day 6, when they had become ‘experts’ (Figure 3C). We recorded the change in Ca^2+^ fluorescence intensity of these neurons as video files, which were then analyzed using the HOTARU^27^ system. This analysis yielded the original data matrix V, which consists of two dimensions: neurons (N) and time (T). The Ca^2+^ activity, represented by fluorescence intensity, for the *n*^th^ neuron at time *t* was recorded in the *n*^th^ row and *t*^th^ column of this matrix.

**Figure 3.**
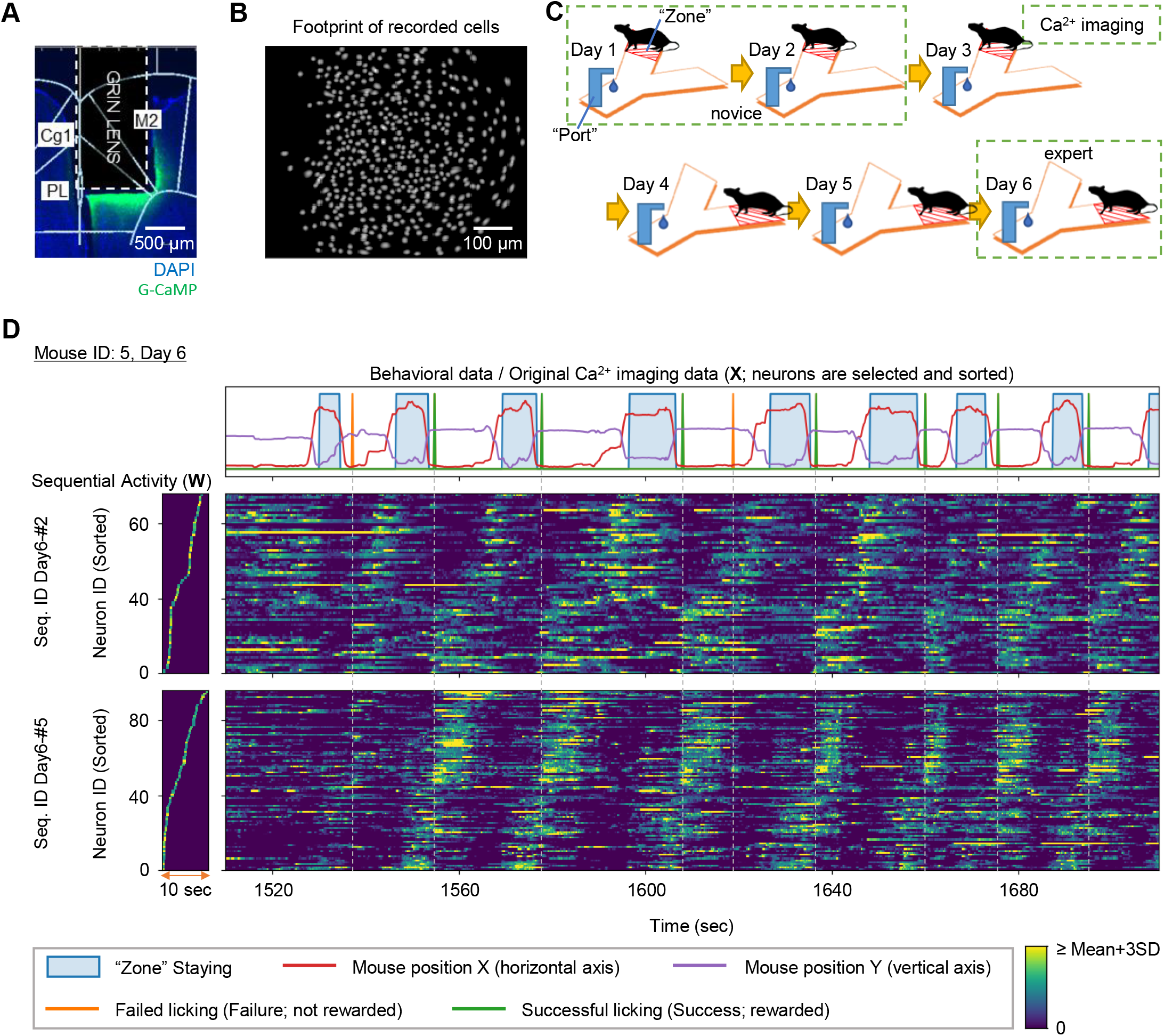
The medial prefrontal cortex (mPFC) contains neuronal sequences that reflect reward acquisition. (A) Location of lens implantation and expression of DAPI (blue) and G-CaMP (green). (B) Footprints of cells recorded through Ca^2+^ imaging. (C) Imaging schedule during Y-maze task learning. (D) Example of neuronal sequences detected on day 6 in mouse ID: 5, representing the behavior of the mouse (top) and the shape of the neuronal sequences (bottom left). For clarity, the displayed neuronal sequences underwent the sequence-sharpening operation (see STAR Methods). Original data sorted according to the order of the neuronal sequences (bottom right). See also Figure S5 and Tables S1 and S2. Abbreviations: Cg1, cingulate area1; PL, prelimbic cortex; M2, secondary motor cortex; Seq., Neuronal sequence.

We then employed iSeq to decompose the original matrix V, collected every 3 days from six mice, into two matrices: pattern matrix W, which preserves the shape of neuronal sequences, and intensity matrix H, which captures the time series of sequential activity values. From these decompositions, we identified 4–10 neuronal sequences, each lasting up to 10 s (Tables S1 and S2 and Figures S5A–S5B). The durations of the detected neuronal sequences and the proportions of their constitutive cells did not vary across experimental days (Figures S5C–S5D). Each neuronal sequence was systematically discriminated by the mouse ID, date, and sequence ID (e.g., Seq. ID: Day1-#2 of mouse 3). Based on data from the pattern matrix W, we extracted cells comprising neuronal sequences from the original matrix V and analyzed their correlation with mouse behavior over time (Figure 3D). We thus discovered neuronal sequences active both before and after reward acquisition. Specifically, some neuronal sequences remained active from the moment the mouse obtained a reward (or licked the Port) until it entered the Zone (e.g., Figure 3D; Seq. ID: Day6-#2). Additionally, other neuronal sequences were activated in anticipation of a reward (e.g., Figure 3D; Seq. ID: Day6-#5). Importantly, iSeq extracts neuronal sequences without referring to behavioral data, and thus the fact that neuronal sequences (shown in Figure 3D) emerged in correspondence with behavior is particularly meaningful. Furthermore, although the two neuronal sequences depicted in Figure 3D sometimes behave as if they are part of a single large sequence, their independent activities can be distinguished, such as by their distinct responses to a Failure lick at approximately 1,530 s.

### Neuronal sequences in the mPFC encode behaviors related to the procedural rule

Based on the findings from Figure 3D, we hypothesized that individual neuronal sequences identified in the mPFC of mice correspond to specific behaviors required to complete the Y-maze task. To precisely assess the timing of neuronal sequence activation, we implemented the sequence-sharpening process (STAR Methods) on the pattern matrix W and intensity matrix H, which encapsulate individual neuronal sequences. Owing to the inherent nature of iSeq and ConvNMF, the activity of a single neuronal sequence can be decomposed into W and H in multiple ways (Figures S6A–S6C). Among these decomposition forms, we modified and standardized the values of W and H to depict neuronal sequences as “narrow” sequences in W (Figure S6A, Figures S6D–S6E). Moreover, to emphasize the onset timing of neuronal sequence activity over its duration, we applied a high-pass filtering technique (commonly used to estimate neuronal spike activity from Ca^2+^ traces^28,29^) to the intensity matrix H. To prevent potential phase shifts due to filtering, we extracted the low-frequency components as a moving average (Figure S6F). By subtracting this moving average from the original activity values and scaling based on the moving average, we derived a measure of spike-like activity referred to as the spike intensity of the neuronal sequences (Figure S6G).

Using data from mouse location tracking, we calculated the average spike intensity for each mouse position (Figure 4A; top). Additionally, we time-shifted the spike intensity data relative to the location data to estimate the movement trajectories of the mouse during the occurrence of neuronal sequences (Figure 4A; bottom).

**Figure 4.**
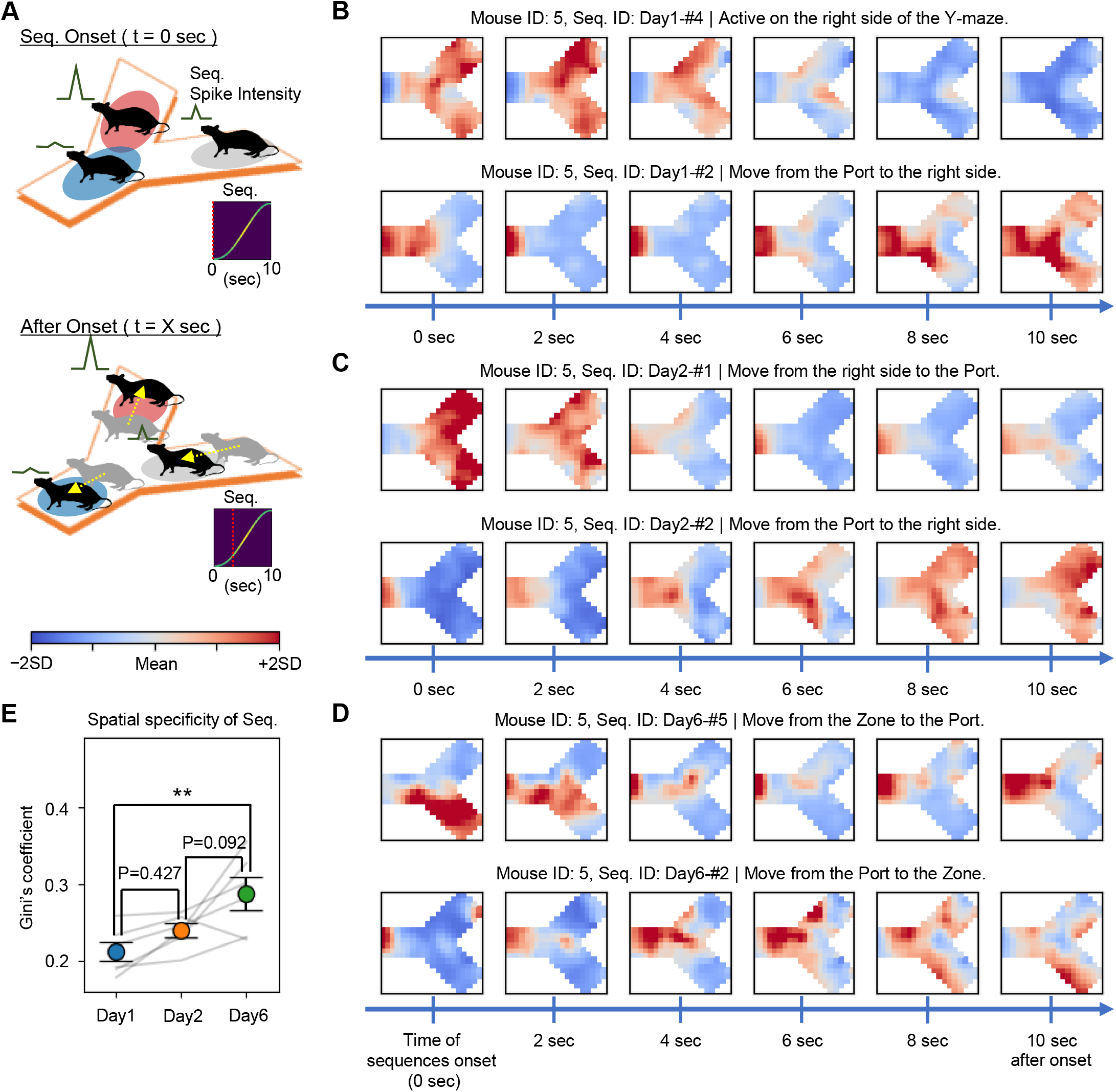
Neuronal sequences in the medial prefrontal cortex (mPFC) encode behaviors related to the rules. (A) Schematic for estimating mouse movement trajectories from the mouse’s location and average spike intensity of each neuronal sequence. (B–D) Examples of estimated mouse movement trajectories of mouse ID: 5 during the activity periods of each neuronal sequence: (B) Day 1, (C) Day 2, and (D) Day 6. Regardless of the duration of each neuronal sequence, the movement trajectories of the mouse were estimated and plotted for 10 s following the onset of the neuronal sequences. (E) The spatial specificity of the neuronal sequences, determined by the average Gini coefficient of the distribution of mouse positions within the estimated trajectories (\*\**P*=0.008; one-way RM ANOVA with Tukey’s post hoc test). See also Figures S6 and S7. Abbreviations: Seq., Neuronal sequence. Data are represented as mean ± SEM.

We compiled these values into what we refer to as a “location vector”, which we displayed as a heatmap formatted to match the shape of the Y-maze (Figures 4B–4D). Our findings indicate that on all days, neuronal sequences correspond to behaviors necessary for completing the rules of the Y-maze task. Specifically, we observed neuronal sequences during movements from the Zone located at the right side to the Port to receive a reward (Figures 4B–4D; upper panels) and from the Port back to the Zone or the right side to obtain the next reward (Figures 4B–4D; lower panels). Interestingly, in mice performing the Y-maze task rule as a three-step procedure (i.e., moving to the opposite arm of the Zone (hereafter called “the Opposite”), from the Opposite to the Zone, and from the Zone to the Port), neuronal sequences corresponding to each of these three behavioral stages were detected (Figures S7A–S7C). By calculating the Gini’s coefficient for the location vector of each neuronal sequence, we could evaluate the spatial specificity of the neuronal sequences, assessing their location bias within the Y-maze. Our results clearly demonstrate that as rule learning progresses, the trajectories of behavior encoded by these neuronal sequences become increasingly distinct (Figure 4E).

### The neuronal sequences of expert mice distinguish between successful and unsuccessful reward acquisition

If neuronal sequences in the mPFC encode procedural rule-related information, the activity patterns of these sequences should vary between successful (Success) and unsuccessful (Failure) reward acquisition (Figure 5A). We calculated the average of the spike intensity for neuronal sequences 20 s before and after licking the Port in both Success and Failure cases (Figure 5B). To assess the differences in neuronal sequence activity patterns between Success and Failure, we analyzed their Pearson’s correlation. The correlation after the mouse licked the Port (post-correlation) decreased compared to before licking the Port (pre-correlation) on all days (Figure 5C), likely due to the difference in outcomes between obtaining a reward in Success cases and not obtaining one in Failure cases. Of note, on day 6, the pre-correlation significantly decreased compared to days 1 and 2 (Figure 5D). This outcome was also replicated when analyzing the pre-correlation during the 10 s before the mouse licked the Port (Figure 5F). These findings show that in the mPFC of mice that have mastered the rule, individual neuronal sequences exhibit distinct activity patterns capable of differentiating between successful and unsuccessful task completions, even before the reward is actually obtained. Additionally, when similar analyses were performed on the post-correlations, the results demonstrated variability depending on the time duration over which the correlation was assessed (Figures 5E and 5G).

**Figure 5.**
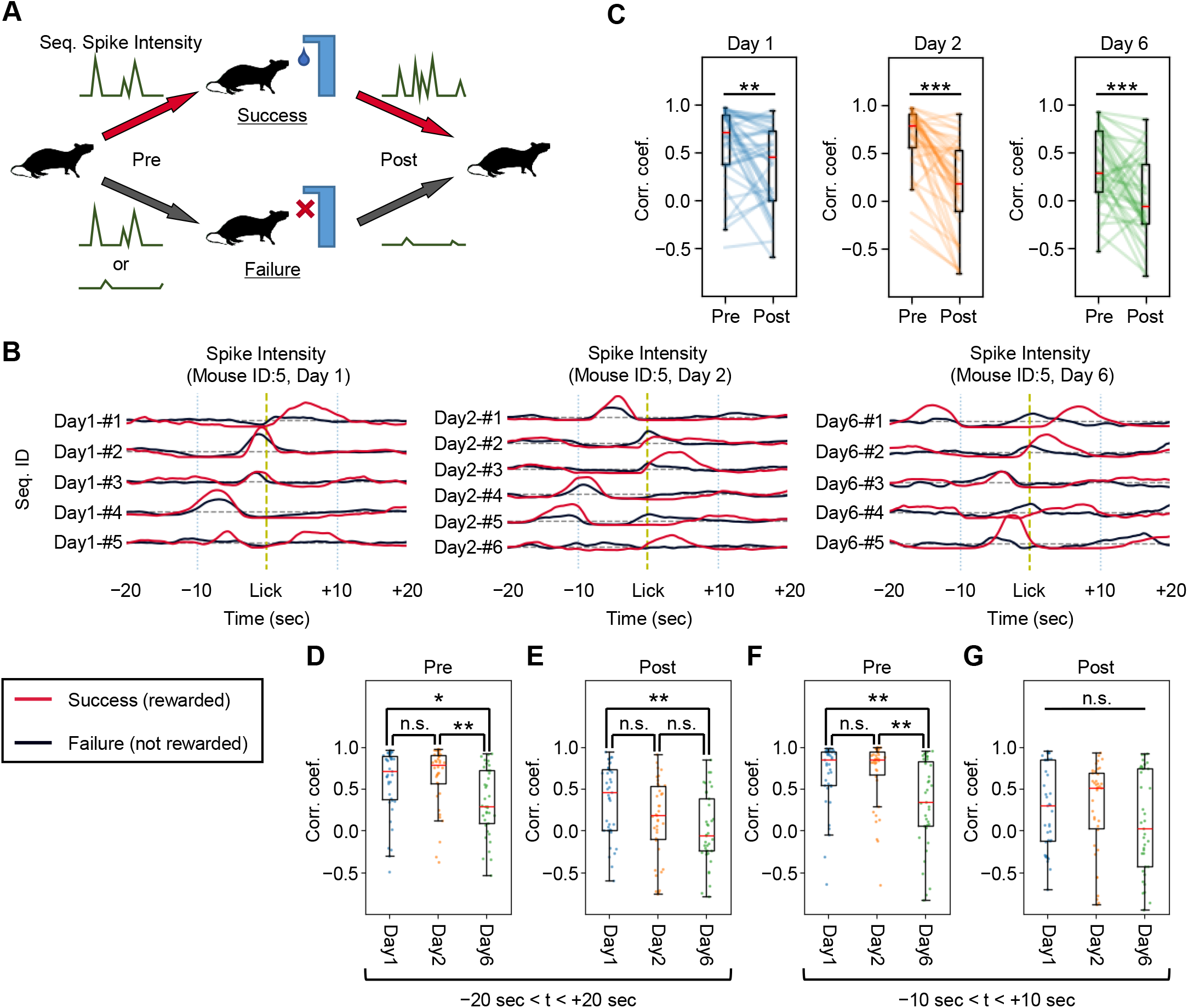
The neuronal sequences of expert mice distinguish between successful and unsuccessful reward acquisition. (A) Schematic for analyzing the difference in activity patterns of neuronal sequences during Success and Failure. (B) Time series of the average spike intensities of each neuronal sequence during the 20 s periods before and after the mouse licked the Port, plotted separately for Success and Failure. (C) Changes in the correlations of spike intensities for Success and Failure before (Pre) and after (Post) the mouse licked the Port (Day 1, \*\**P*=0.003; Day 2, \*\*\**P*<0.001; Day 6, \*\*\**P*<0.001; t-test on two related samples of scores). (D) Daily comparison of spike intensity correlations in the pre-lick phase, between Success and Failure (\**P*=0.016; \*\**P*=0.003; one-way ANOVA with Tukey’s post hoc test). (E) Same as (D), but in the post-lick phase (\*\**P*=0.002). (F) Same as (D), but focused on the 10 s prior to licking (Days 1–6, \*\**P*=0.003; Days 2–6, \*\**P*=0.001). (G) Same as (E), but focused on the 10 s after licking. Abbreviations: Seq., Neuronal sequence; Corr. Coef., Correlation coefficient. Data are represented as mean ± SEM.

### The dynamics of neuronal sequences of expert mice distinguish between successful and unsuccessful reward acquisition

Furthermore, we explored whether there were differences in the overall neuronal dynamics driven by multiple neuronal sequences, rather than just individual ones, between novice and expert mice. We employed principal component analysis (PCA) to extract the first and second principal components (Figure 6A), allowing us to represent the spike intensity of all detected neuronal sequences from each mouse on each day in a two-dimensional space (Figure 6B). We then plotted the average trajectories of these principal components during the 20 s before licking the Port, in both Success and Failure cases, defining these plotted trajectories as sequential neuronal dynamics. Our analysis focused on determining whether these dynamics were similar or differed between Success and Failure cases (Figure 6C). The results indicate that the sequential neuronal dynamics on days 1 and 2 were similar for both outcomes, whereas those on day 6 differed considerably between Success and Failure (Figure 6D). By applying the same analytical methods as used in Figure 2G, we observed that the distance between the dynamics of expert mice was significantly greater (Figures 6E–6F). Given the consistent differences in the behavioral trajectories of mice between Success and Failure across all days (Figure 2G), the significant distinction in the sequential neuronal dynamics of expert mice most likely does not reflect changes in external stimuli, such as location shifts due to altered behavior. Instead, it indicates that these differences stem from the mouse’s ability to distinguish between Success and Failure, a result of their understanding of the rule.

**Figure 6.**
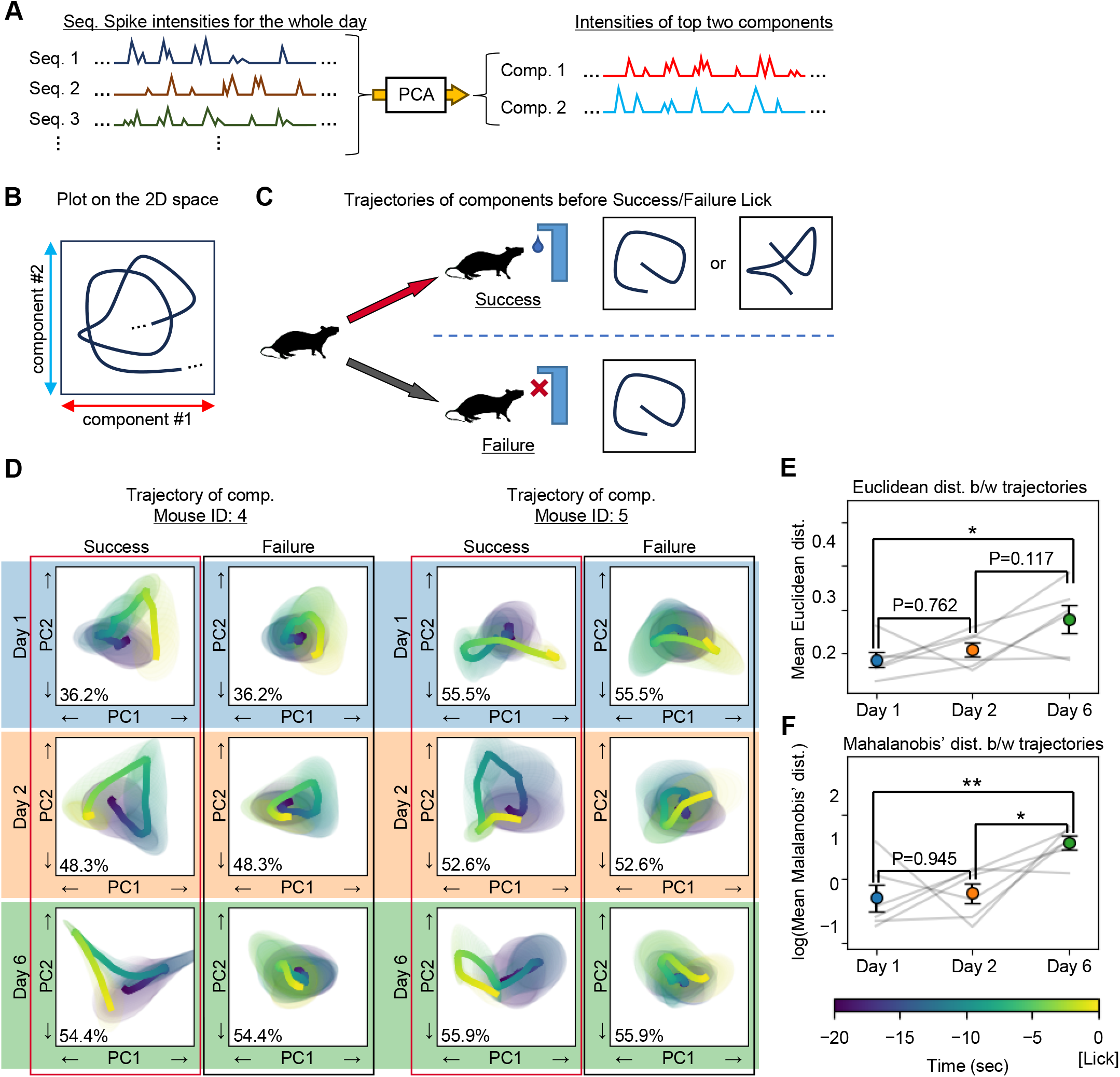
The dynamics of neuronal sequences of expert mice distinguish between successful and unsuccessful reward acquisition. (A) Schematic for converting the time series of spike intensities from neuronal sequences into the top two principal components of sequential neuronal dynamics. (B) Schematic for plotting these sequential neuronal dynamics in a two-dimensional space. (C) Schematic for assessing whether distinct sequential neuronal dynamics occur in Success and Failure. (D) Plots of the sequential neuronal dynamics for mice ID: 4 and 5 during the 10 s period leading up to Success and Failure. The lines indicate the average trajectories of these dynamics. The shaded areas around the lines represent probability ellipses, statistically depicting the regions through which 50% of the sequential neuronal dynamics pass. The inset (bottom left of each trajectory) shows the sum of the explained variance ratios for the top two principal components. (E–F) The differences in sequential neuronal dynamics between Success and Failure were assessed by the same method as employed in Figure 2G. (E) Euclidean distance (\**P*=0.031; one-way RM ANOVA with Tukey’s post hoc test). (F) Mahalanobis’ distance (\*\**P*=0.006; \**P*=0.011; one-way RM ANOVA with Tukey’s post hoc test). Abbreviations: Seq., Neuronal sequence; Comp., Component; PC, Principal component; Dist., Distance; b/w, Between. Data are represented as mean ± SEM.

### In expert mice, neuronal sequences show improved decoding accuracy only for behaviors crucial to the procedural rule

If neuronal sequences encode procedural rule-related information, it should be possible to decode mouse behaviors from the spike intensities of these sequences. We implemented a decoder featuring an LSTM (long short-term memory) layer^30^ to decode mouse behaviors from the temporal activity patterns of neuronal sequences (Figure 7A). We assessed the decoder’s accuracy across various proficiency levels of the mice. To prevent overfitting, the data were split into Training and Validation datasets (Figure 7B). Specifically, we used only the Training data to train the decoder and terminated training when the prediction accuracy on the Validation data reached its minimum (Figure 7C; see also STAR Methods).

**Figure 7.**
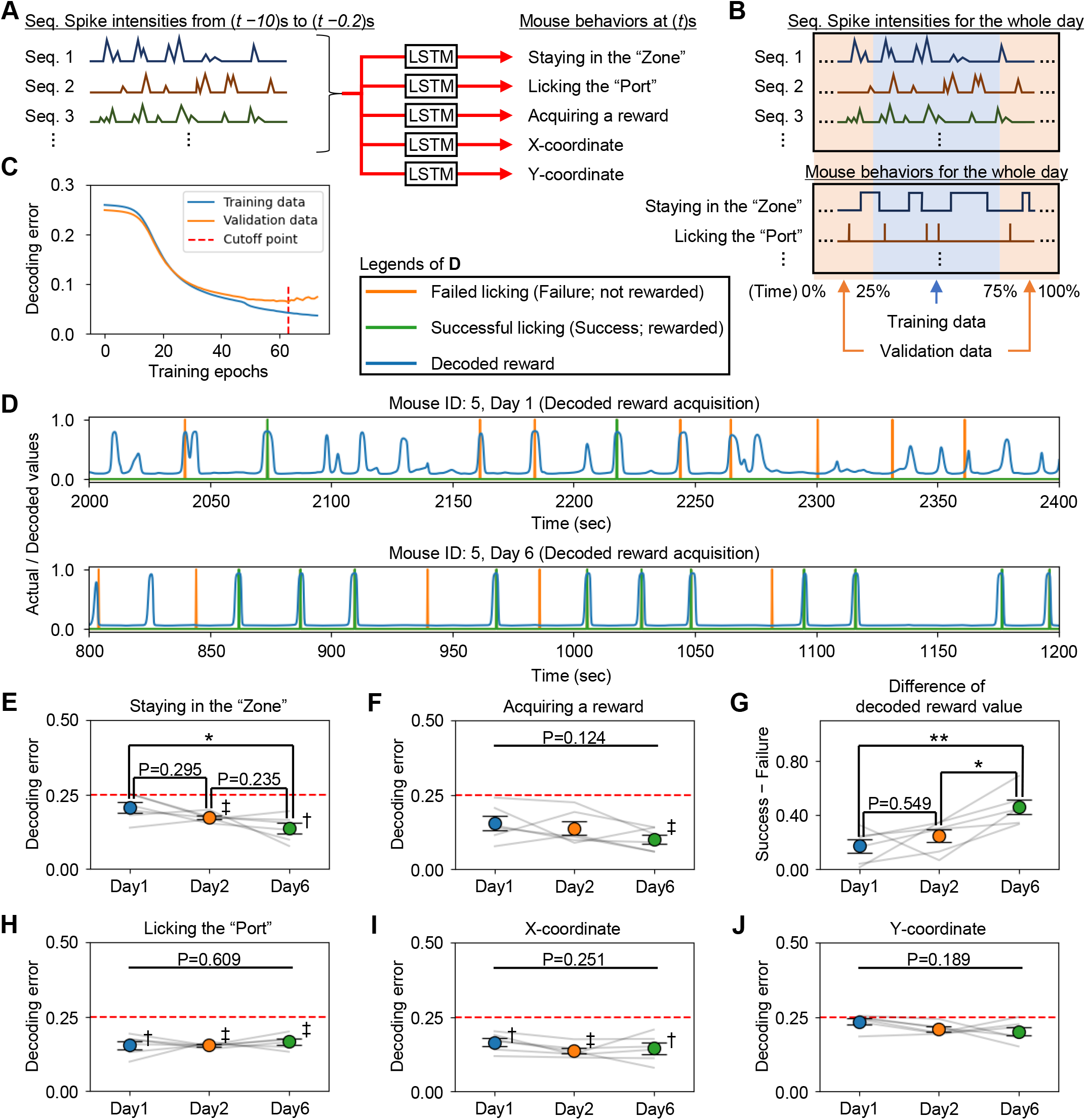
In expert mice, neuronal sequences show improved decoding accuracy only for behaviors crucial to the rule. (A) Schematic for decoding mouse behavior from the time series of spike intensities in neuronal sequences. (B) Schematic for dividing data into Training and Validation datasets. (C) Example of prediction error fluctuations for Training and Validation data, with training of the decoder halted at the epoch marked by the red dashed line, designated as the final result. (D) Reward values predicted by the decoder, displayed alongside the actual instances of Success and Failure for the mouse. (E) Daily decoding errors for “staying in the Zone” (\**P*=0.014; one-way RM ANOVA with Tukey’s post hoc test). The red dashed line indicates the baseline error from the null decoder (Day 1, *P*=0.84; Day 2, ‡*P*<0.01; Day 6, †*P*=0.02; comparison with the null decoder using a one-sample t-test with Bonferroni’s correction). (F) Same as (E), but for “acquiring a reward” (Day 1, *P*=0.17; Day 2, *P*=0.07; Day 6, ‡*P*<0.01). (G) Differences in the outputs of the decoder predicting “acquiring a reward” at the times of Success and Failure licking (\*\**P*=0.003; \**P*=0.023 one-way RM ANOVA with Tukey’s post hoc test). (H) Same as (E), but for “licking the Port” (†*P*=0.02; ‡*P*<0.01). (I) Same as (E), but for the “X-coordinate” (Day 1, †*P*=0.02; Day 2, ‡*P*<0.01; Day 6, †*P*=0.04). (J) Same as (E), but for the “Y-coordinate” (Day 1, *P*=1.00; Day 2, *P*=0.05; Day 6, *P*=0.31). Abbreviations: Seq., Neuronal sequence; LSTM, Long short-time memory. Data are represented as mean ± SEM.

The decoding accuracy for critical behaviors in the Y-maze task (“staying in the Zone” and “acquiring a reward”) significantly improved on day 6 compared to days 1 and 2 (Figures 7D–7G). Although the overall average decoding accuracy for “acquiring a reward” did not exhibit statistically significant changes across the days, it was only on day 6 that the accuracy significantly outperformed the null decoder (Figure 7F; see also STAR Methods). Additionally, when analyzing the decoder’s output exclusively at the moments of licking, the results from day 6 clearly demonstrated the decoder’s ability to distinguish between Success and Failure (Figure 7G). These results indicate that as mice learn the rule, key behaviors crucial to the rule are progressively encoded into the neuronal sequences.

Conversely, the decoding accuracy for “licking the Port”, which does not differentiate between Success and Failure, was significantly high from day 1 and remained consistent regardless of the mice’s proficiency in the task (Figure 7H). Although mice started learning the rules of the Y-maze task from day 1, port habituation had been initiated 2 days earlier, enabling the mice to recognize the meaning of the Port by day 1 (see STAR Methods). Given this context, Figure 7H indicates that information regarding the Port was already encoded in the neuronal sequences by day 1.

Regarding the decoding accuracy for X- and Y-coordinates, no significant changes were observed across different days. Of note, the decoding accuracy for the X-coordinate, which strongly reflects the position of the Port, consistently surpassed that of the null decoder (Figure 7I). However, the decoding accuracy for the Y-coordinate did not differ from that of the null decoder (Figure 7J). These findings, in conjunction with those displayed in Figures 7D–7G, indicate that the neuronal sequences are encoding information not only based on location but rather on the value of behaviors associated with the rule.

### The populations of cells of the neuronal sequences are continuously updated

The cells constituting the neuronal sequences appeared to be dispersed across the field of view used for Ca^2+^ imaging (Figure 8A). Employing a method similar to that used by Harvey et al.^15^, we classified cell pairs into six groups based on their distances and analyzed the differences in their activity levels within a single neuronal sequence (Figure 8B). Although the trend was gradual, there was a tendency for cells that were closer together to less frequently be part of the same neuronal sequence. This pattern suggests the involvement of local inhibitory neurons in the formation of neuronal sequences.

**Figure 8.**
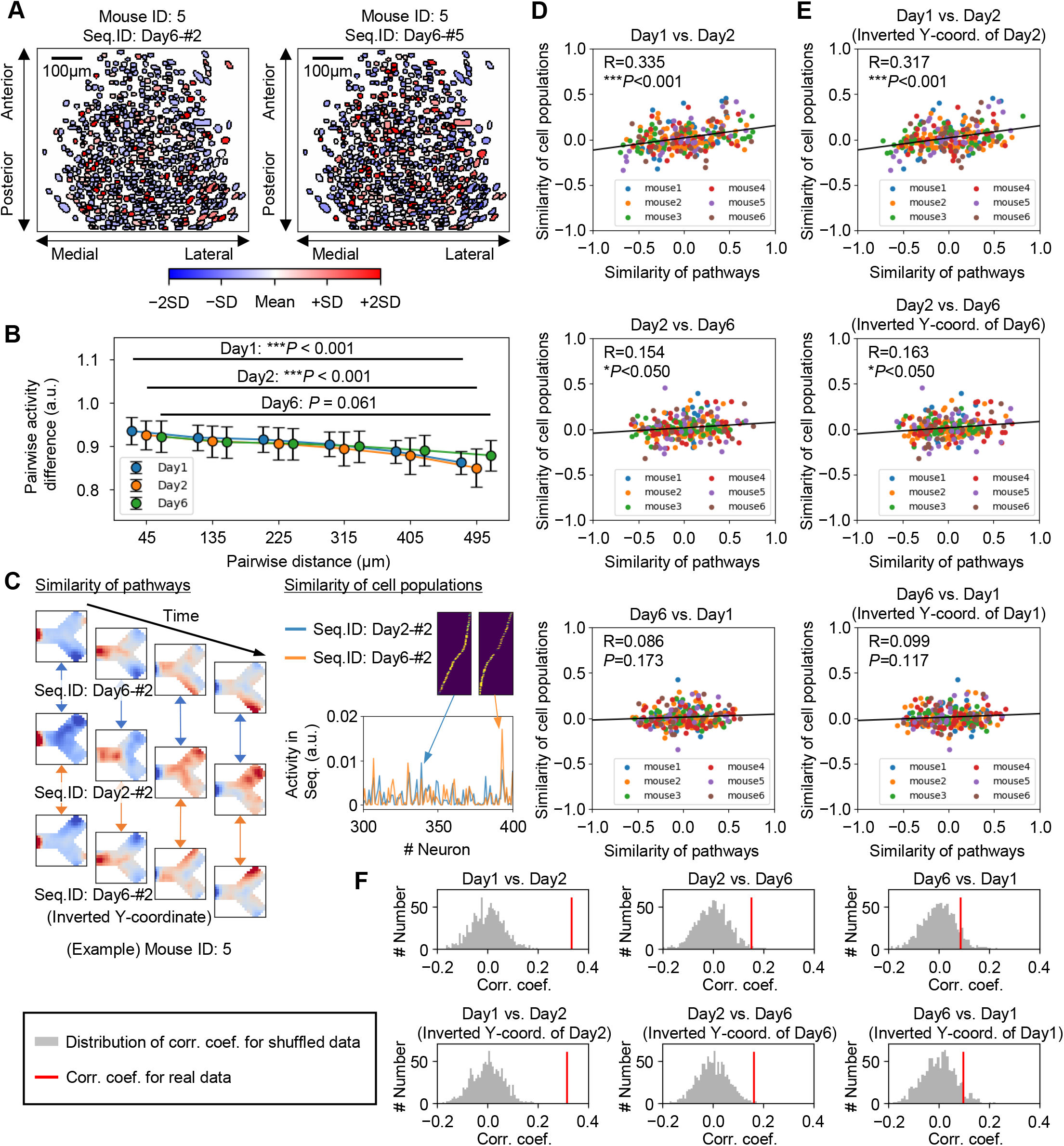
The populations of cells of the neuronal sequences are continuously updated. (A) An example illustrating the activity intensity of cells within a single neuronal sequence and their spatial distribution. (B) The relationship between the spatial distance of two cells and the differences in their activity intensities within a neuronal sequence (Friedman’s test for repeated samples with Bonferroni’s correction). (C) Schematic showing the similarity of pathways encoded by neuronal sequences and the similarity among the cell populations composing these sequences. (D) Correlation between the similarity of pathways and the similarity of cell populations in neuronal sequences. The correlation coefficient R and *P* value (Wald’s test with t-distribution) are shown. (E) Same as (D), but with inverted pathway Y-coordinates on one day. (F) Comparison of the correlation coefficients obtained in (D) and (E) with the distribution from 1000 simulations in which cell IDs were shuffled and recalculated. Abbreviations: Seq., Neuronal sequence; Corr. Coef., Correlation coefficient; Coord., Coordinate. Data are represented as mean ± SEM.

Finally, we investigated the day-to-day consistency of the cells of neuronal sequences. Neuronal sequences of novice mice were already associated with specific behaviors (Figure 4). However, as the mice progressed in learning, the behaviors associated with the rule became more precisely encoded within the neuronal sequences. Thus, even if certain neuronal sequences during the novice phase corresponded to behaviors similar to those in the expert phase, these sequences would likely undergo modifications as the mice’s understanding of the rules deepened. We evaluated the similarity of behaviors encoded in neuronal sequences by analyzing the similarity of mouse trajectories during the activity of these sequences (Figure 4). Additionally, we examined the correlation with the similarity of the cells of the neuronal sequences (Figure 8C). Although there was a modest consistency in the sequences between days 1 and 2, there was little overlap between the sequences of novice and expert mice (Figure 8D). Considering the reversal of the Zone’s location on day 6 compared with days 1 and 2, we also computed the correlation after inverting the Y-axes of the trajectories of one day; however, the trend did not change (Figure 8E–8F). These results indicate that the cells forming neuronal sequences are being updated daily.

## Discussion

Our findings highlight the critical role of neuronal sequences in the PFC. Not only do individual neuronal sequences encode specific behaviors (Figures 4 and S7) but collectively, multiple sequences create dynamics that represent the procedural rule, enabling the PFC to differentiate between processes that lead to successful and unsuccessful reward acquisition (Figures 5 and 6). As animals advance in procedural rule learning, the dynamics of their neuronal sequences increasingly concentrate on encoding behaviors essential to the rule (Figure 7). This indicates that neuronal sequences are continuously restructured each day throughout this process (Figure 8). It is likely that research focusing on neuronal sequences will continue to be crucial, and the iSeq algorithm we used will prove highly valuable in this context. As neuroimaging technologies advance, the volume of data generated from experiments is rapidly increasing. However, iSeq can automatically detect neuronal sequences within this data, significantly decreasing the analytical workload.

In this study, a consistent and significant finding is that the neuronal sequences in the mPFC encode the meaning of the mouse’s behaviors more strongly than merely the positional changes resulting from those behaviors. For instance, although no significant changes were observed in the differences between the movement trajectories leading to Success and Failure as a result of procedural rule learning (Figure 2G), the trajectories of sequential neuronal dynamics in the mPFC of expert mice clearly distinguished between these outcomes (Figures 6D–6F). Additionally, even as animals learned the rule, the decoding accuracy for the X-or Y-coordinates based on sequential neuronal dynamics did not improve (Figures 7I–7J). In particular, the fact that the decoding accuracy for Y-coordinates consistently matched that of the null decoder suggests that the sequential neuronal dynamics in the mPFC do not significantly encode information regarding Y-coordinates. Conversely, the decoding accuracy for X-coordinates consistently and significantly exceeded that of the null decoder. This is likely because the X-coordinates are strongly linked to the location of the Port.

A previous study has shown that although hippocampal neurons exhibit activity characteristics that respond to the location of objects, neurons in the PFC exhibit equivalent responses to objects with the same meaning, regardless of their location^31^. Furthermore, hippocampal neurons encode absolute positions, whereas neurons in the PFC encode positions relative to the animal’s perspective^4,18^. This supports the model suggesting that the PFC converts sensory inputs into specific behavioral outputs^32-34^. Indeed, in the PFC of macaque monkeys, only sensory inputs critical to the task are integrated into the neuronal population dynamics specific to their choices^32^. Therefore, it is more likely that the meanings of objects (rather than their identities) and the relative positions of a mouse (rather than its absolute positions), are incorporated into the populational representation of PFC neurons. In our experiments, whether the mouse was in the Zone or not was more crucial to the task than its exact X-or Y-coordinates, and thus it is conjectured that the dynamics of the neuronal sequences evolved to encode the information of the Zone as the mice learned the rule. These results strongly support our hypothesis that the PFC recognizes the meaning of stimuli and actions, selects elements crucial for reward acquisition, encodes them into neuronal sequences, and thereby represents the procedural rule.

Figures 4 and S7 display the behaviors encoded by individual neuronal sequences. Specific neuronal sequences were active when mice performed actions corresponding to their understanding of the rule (mouse ID: 5, two steps; mouse ID: 6, three steps). These findings may suggest that neuronal sequences decompose the rule into multiple actions for encoding. This aligns with the hypothesis of “compositionality”^35-38^ of the PFC, i.e., remembering complex tasks as sets of individual components enables rapid learning when faced with new tasks that share common elements. In this study, we exposed mice to only one rule; nevertheless, introducing multiple similar rules simultaneously in future experiments could enable the identification of common neuronal sequences that span across various rules.

Our results show that neuronal sequences and their dynamics encode procedural rules; however, the underlying mechanisms generating these sequential activities and their effects remain unclear. A possible hypothesis is that the mPFC orchestrates procedural rule execution through the activity of neurons with diverse connectivities. The PFC is interconnected with multiple brain regions^39^. In the mPFC, sequentially activating neurons that connect to regions such as the sensory cortex (sensation), amygdala (emotion), hippocampus (memory), and motor cortex (action) could integrate processes across these areas. This integration might enable the animal to evaluate external stimuli and execute appropriate actions based on memory, thereby facilitating procedural rule-based behavior. To optimize this process, it may be necessary to reconfigure the cell populations of the neuronal sequences, forming mPFC circuits more aligned with this rule. Investigating this mechanism of updating neuronal sequences should be a priority for future research. A previous study has shown that spike-timing-dependent plasticity (STDP) plays a role in the retention^40^ and elimination^41^ of neuronal sequences. Additionally, local inhibitory neurons may affect the formation of these neuronal sequences (Figure 8B).

In conclusion, as procedural rule learning progresses, neuronal sequences in the mPFC encode behaviors critical to the rule, and their dynamics effectively represent this rule. Consequently, the cell populations forming these neuronal sequences are continuously updated to facilitate this process. Future research should aim to elucidate the mechanisms by which neuronal sequence activity triggers procedural rule-based behavior and neuronal sequences are reconfigured to align with rules.

## Resource availability

### Lead contact

Further information and requests for resources and reagents should be directed to and will be fulfilled by the lead contact, Kaoru Inokuchi.

## Materials availability

This study did not generate new unique reagents.

## Data and code availability

- Data reported in this paper is available from the lead contact upon request.
- The Python code used to analyze data in this paper and the iSeq package running on a GPU (igpu) are available at the Github repository (https://github.com/IdlingBrainUT/Ohno_iSeq).
- Any additional information required to reanalyze the data reported in this paper is available from the lead contact upon request.

## Acknowledgements

This work was supported by the JSPS KAKENHI (grant numbers: JP18H05213, JP23H05476), the Core Research for Evolutional Science and Technology (CREST) program (JPMJCR23N2) of the Japan Science and Technology Agency (JST), and the Takeda Science Foundation to K.I.; the AMED PRIME (23gm6510028h0001), the JSPS KAKENHI (20H03554, 23H02785), the Uehara Memorial Foundation, the Naito Foundation, the Inamori Foundation, the Takeda Science Foundation, the Firstbank Of Toyama Scholarship Foundation Research Grant, the Hokuriku Bank Grant-in-Aid for Young Scientists, and the Tamura Science and Technology Foundation to M.N.; and the JSPS KAKENHI (24K18607) support to S.O. S.O. would like to thank Prof. Michikazu Sekine, Department of Epidemiology and Health Policy, University of Toyama, for his long-standing support.

## Author contributions

Conceptualization, S.O. and K.I.; methodology, S.O. and M.N.; formal analysis, S.O.; investigation M.N.; resources, M.N. and K.I.; writing – original draft, S.O; writing – review & editing, M.N. and K.I.; supervision, K.I.; funding acquisition, M.N. and K.I.

## Declaration of generative AI and AI-assisted technologies in the writing process

During the preparation of this work, the authors used ChatGPT to refine English expressions. After using this tool, the authors reviewed and edited the content as needed and take full responsibility for the content of the published article.

## Supplemental information

Document S1. Figures S1–S7, Tables S1–S2, and Math Notes S1–S2.

Video S1. Daily task performance of the mouse during the Y-maze training, related to Figure 2.

## Table

We do not have main tables.

## STAR Methods

### EXPERIMENTAL MODEL AND STUDY PARTICIPANT DETAILS

#### Animals

Naïve wild-type male C57BL/6J mice were purchased from Sankyo Labo Service Co. Inc. and maintained on a 12 h light/dark cycle at a controlled temperature (24□± 3 °C) and humidity (55 ± 5%) with free access to food and water. Mice used in behavioral experiments were 3–5 months old. All the behavioral experiments were performed between zeitgeber ZT+1 and ZT+9. All experimental procedures with animals were approved by the Animal Care and Use Committee of the University of Toyama (approval numbers: A2019MED-35 and A2022MED-7) and were conducted in accordance with the Institutional Animal Experiment Handling Rules of the University of Toyama and the guidelines of the NIH.

## METHOD DETAILS

### Viral vectors

For in vivo Ca^2+^ imaging experiments, the recombinant AAV vector used was AAV_9_-SynI::janelia G-CaMP7b^42^ (9.4□×□10^12^ vg/mL), which was diluted 10-fold in phosphate-buffered saline (PBS; T900, Takara Bio Inc.) before injection. The pAAV-SynI-janelia-G-CaMP7b plasmid, a gift from Douglas Kim (Addgene plasmid #104489), was previously constructed. Recombinant AAV_9_ production was performed using the minimal purification method, and the viral genomic titer was subsequently calculated as described previously^43^.

Briefly, pAAV recombinant vectors were produced using AAV293 cells (240073, Agilent Tech) cultured in 15 cm dishes (Corning). Cultured cells were maintained in Dulbecco’s Modified Eagle Medium (11995-065) supplemented with 10% fetal bovine serum (FBS) (10270106), 1% 2 mM L-glutamine (25030-149), 1% 10 mM nonessential amino acid (MEM NEAA 100×, 11140-050), and 1% (100×) penicillin–streptomycin solution (15140-148, all from GIBCO Life Technologies). AAV293 T cells that reached 70% confluency were transfected using medium containing the constructed expression vector, pRep/Cap, and pHelper (240071, Agilent Technologies), mixed with the transfection reagent polyethylenimine (PEI) hydrochloride (PEI Max, 24765-1, Polysciences Inc.) at a 1:2 ratio (W/V). After 24 h, the transfection medium was discarded, and cells were incubated for another 5 days in a maintenance medium without FBS. On day 6, the AAV-containing medium was collected and purified from cell debris using a 0.45 μm Millex-HV syringe filter (SLHV033RS, Merck Millipore). The filtered medium was concentrated and diluted with D-PBS (14190-144, GIBCO Life Technologies) twice using a Vivaspin 20 column (VS2041, Sartorius) after blocking the column membrane with 1% bovine serum albumin (01862-87, Nacalai Tesque Inc.) in PBS.

To further calculate the titer, degradation of any residual complementary DNA (cDNA) in the viral solution from production was first assured by Benzonase nuclease treatment (70746, Merck Millipore). Subsequently, viral genomic DNA was digested with proteinase K (162-22751, FUJIFILM Wako Pure Chemical) and extracted with phenol/chloroform/isoamyl alcohol (25:24:1 vol/vol), followed by precipitation with isopropanol and final dissolution in Tris-EDTA buffer (10 mM Tris, pH 8.0, 1 mM ethylenedinitrilo tetraacetic acid [EDTA]). The titer quantification for each viral solution, referenced to that of the corresponding expression plasmid, was performed by real-time qPCR using THUNDERBIRD SYBR qPCR Master Mix (QPS-201, Toyobo Co. Ltd.) with the primers 5′-GGAACCCCTAGTGATGGAGTT-3′ and 5′-CGGCCTCAGTGAGCGA-3′ targeting the inverted terminal repeat (ITR) sequence. The cycling parameters were adjusted as follows: initial denaturation at 95 °C for 60 s, followed by 40 cycles of 95 °C for 15 s and 60 °C for 30 s^44^.

### Stereotactic surgery for Ca^2+^ imaging

Mice (3–5 months old) were sedated with an intraperitoneal anesthetic injection^45^ containing 0.75 mg/kg medetomidine (Domitor; Nippon Zenyaku Kogyo Co. Ltd.), 4.0 mg/kg midazolam (Fuji Pharma Co. Ltd.), and 5.0 mg/kg butorphanol (Vetorphale, Meiji Seika Pharma Co. Ltd.) and were then placed on a stereotactic apparatus (Narishige). After surgery, an intramuscular injection of 1.5□mg/kg atipamezole (Antisedan; Nippon Zenyaku Kogyo Co.), an antagonist of medetomidine, was administered to boost recovery from sedation. Mice were allowed to recover from surgery for 3 weeks in their home cages before behavioral experiments were initiated.

Viral injections were made using a 10 µL Hamilton syringe (80030, Hamilton) fitted with a mineral oil-filled glass needle and wired to an IMS-20 automated motorized microinjector (Narishige). Mice were injected with 500 nL of AAV_9_-SynI::janelia G-CaMP7 at 100 nL/min unilaterally into the right mPFC (−2.0 mm AP, +0.3 mm ML, +2.2 mm DV). After 2 weeks of recovery from AAV injection surgery, the reanesthetized mice were again placed on a stereotactic apparatus to implant a gradient index (GRIN) lens^45^ (1.0□mm diameter, 4.0□mm length; Inscopix Inc.) into the center of the injection (from the skull surface: +1.2 mm DV) using custom-made forceps attached to a manipulator (Narishige). Low-temperature cautery was used to emulsify bone wax into the gaps between the GRIN lens and the skull, and then the lens was anchored in place using dental cement. Three weeks after GRIN implantation, the mice were reanesthetized and placed back onto the stereotactic apparatus to set a baseplate (Inscopix Inc.), as described previously^45^. In brief, a Gripper (Inscopix Inc.) holding a baseplate attached to a miniature microscope (nVista 3; Inscopix Inc.) was lowered over the previously set GRIN lens until visualization of clear vasculature was possible, indicating the optimal focal plane. Dental cement was then applied to fix the baseplate in position to preserve the optimal focal plane. Mice recovered from surgery in their home cages for 1 week before the initiation of behavioral imaging experiments.

### Y-maze task

All behavioral experiments with or without Ca^2+^ imaging were performed under dim light conditions (approximately 2 lx). The behavioral apparatus, including a solenoid valve for the regulation of sucrose water delivery (LHDA2431115H, The Lee Company), video tracking sensors (VTS4, BRC), a licking port (Mouse Behavior Port, Sanworks), and white LED tape (SOF-18W-30-SMD, Akizuki Denshi Tsusho), was controlled. Outputs were recorded by in-house-built signal regulators and the OpenEx Software Suite (RX8-2, Tucker-Davis Technologies). OpenEx was also used to synchronize the onsets of behavioral and imaging systems.

The Y-maze context consisted of a trunk and upper and lower branching arms (wall height: 25.5 cm, path width: 12.0 cm, trunk length: 17.0 cm, branching arm length: 16.5 cm). There was a licking port (“Port”) at the end of the trunk, and an area of up to 11.5 cm from the end of either branching arm was defined as the reward chance waiting area (“Zone”). Each end was marked with a different wall pattern (a black dot or stripe). For port habituation, mice were subjected to water deprivation in their home cage from 1 day before the port habituation and Y-maze training sessions and were then rehydrated with a water bottle for 1 h after each behavioral session. Before the Y-maze training, mice were individually subjected to port habituation sessions for 2 days, in which the mice were habituated to licking the Port to drink 10% sucrose water in a small cage until they had drunk approximately 1.5 mL. The water supply from the tap of the Port was controlled so that it only delivered one 7.5 μL water drop when the animal licked the Port.

For the Y-maze training, the animal was required to learn three behaviors to obtain reward water in the Y-maze context. First, the animal needed to stay in the Zone of the arm for the appropriate time until a reward chance emerged. Second, during the 10 s reward chance, the animal needed to move to the Port. Third, the animal needed to lick the Port to drink a 25 μL water drop within the reward chance time. Otherwise, the animal failed to obtain the reward because the reward chance was not induced. The Y-maze training was composed of sequential 6-day trainings. The Zone position and time for inducing the reward chance were altered according to the day of the training session: on day 1, upper Zone for 4 s; on days 2–3, upper Zone for 5 s; on day 4, lower Zone for 5 s; on days 5–6, lower zone for 6 s.

During the training, mice were allowed to freely explore within the Y-maze context and the Port. When mice stayed in the Zone of the Y-maze arm, LED blinking (50 ms pulse, 1 Hz) started (reward chance induction phase) and continued until the reward chance was induced, provided the mice remained in the Zone. However, if the mice moved out of the Zone, the LED blinking stopped. If mice stayed within the Zone for the appropriate time, the reward chance with continuous LED lighting followed the LED blinking. Thus, during the initial learning, the animal unintentionally obtained reward water when it showed the sequential three behaviors: staying in the Zone until a reward chance emerged, moving to the lick port arm, and licking the Port within the reward chance. Mice were subjected to the Y-maze task until they acquired approximately 40 rewards or until 60 min had elapsed. The animal’s XY positions were extracted by the MATLAB program OptiMouse^46^ (https://github.com/yorambenshaul/optimouse) from screen captures from the recorded movies using behavioral movie software (AG-Desktop recorder, AmuseGraphics). Ca^2+^ imaging was performed during the task only on days 1, 2, and 6 to avoid photo bleaching and phototoxicity owing to the long-exposure imaging.

### Behavioral data processing

We developed image processing software in Python to adjust and align the differences in the numbers of sampled frames due to the differences in core clock frequency between the RX8-2 (Tucker-Davis Technologies) at 1017 Hz, nVista (Inscopix) at 1000 Hz, and screen capturing software (AG-Desktop recorder, AmuseGraphics) of the PC at 20.004 Hz. This procedure was performed by referring to the captured video of the computer screen during the experiment.

After one lick of the Port, the mice continued to exhibit several licking behaviors, regardless of whether they were rewarded. This behavior was thought to be a retrieval of the remaining water or a confirmation that the task had really failed when the reward was not obtained. Therefore, we did not count this “just in case” lick as a failed (non-rewarded) lick. That is, until 10 s had elapsed after one lick, if a subsequent lick was not accompanied by a reward, we removed the subsequent lick from the behavioral data.

### In vivo Ca^2+^ imaging data acquisition and analysis

Ca^2+^ signals produced from G-CaMP7 protein expressed in mPFC neurons were captured at 20 Hz with nVista acquisition software (IDAS, Inscopix) at the maximum gain and optimal power of the nVista LED. Ca^2+^ imaging movies were then extracted from the nVista Data acquisition box (DAQ, Inscopix). Using Inscopix Data Processing Software (IDPS, Inscopix), movies were temporally stitched together to create a full movie that contained recordings of all sessions across all days, which was spatially down-sampled (4×) and then corrected for across-day motion artifacts against a reference frame that was chosen from any day that had a clear landmark vasculature. Further motion correction was then applied using Inscopix Mosaic software, as previously described^45^. The full movie was then temporally divided into the individual days using Inscopix Mosaic software. Each movie of individual days was then low-band-pass filtered using Fiji software^47^ (NIH) to reduce noise, as described previously^23,45^. The fluorescence signal intensity change (Δ*F*/*F*) for each day was subsequently calculated using Inscopix Mosaic software according to the formula Δ*F*/*F = (*Δ*F*/*F*_*m*_*)/ F*_*m*_, where *F* represents each frame’s fluorescence, and *F*_m_ represents the mean fluorescence of the whole day’s movie^23,45^. Afterwards, movies representing each day were again concatenated to generate the full movie for all days in the Δ*F*/*F* format. Finally, cells were identified using a HOTARU automatic sorting system^48^, and each cell’s Ca^2+^ signals over time were extracted in a (Ď; time × neuron) matrix format, as previously described^23,45^. This matrix was then transposed and treated as the original matrix V.

Binning was applied every four time frames (0.2 s) to reduce the computational complexity in iSeq, which is described later. In addition, the signal intensity of the cells was scaled so that the average of the nonzero signal intensity of each cell was 1. The maximum signal intensity was capped to be within three standard deviations from the mean of the nonzero signal intensities.

## Immunohistochemistry and microscopy

Immunohistochemistry was conducted as described previously^49^. Mice were deeply anesthetized with a mixed anesthesia solution, as described above, and perfused transcardially with 4% paraformaldehyde in PBS (pH 7.4). Their brains were removed and further post-fixed by immersion in 4% paraformaldehyde in PBS for 24 h at 4 °C. Each brain was equilibrated in 25% sucrose in PBS for 2 days and then frozen in dry-ice powder. Coronal sections of 50 μm thickness were cut on a cryostat and stored at −20 °C in cryoprotectant solution^49^ (25% glycerol, 30% ethylene glycol, and 45% PBS) until further use. For immunostaining, sections were transferred to 12-well cell culture plates (Corning, USA) containing Tris-buffered saline (TBS-T) buffer (0.2% Triton X-100 and 0.05% Tween-20).

For G-CaMP detection, after washing with TBS-T buffer, the floating sections were treated with blocking buffer (5% normal donkey serum [S30, Chemicon, USA] in TBS-T) at room temperature for 1 h. Reactions with primary antibodies were performed in blocking buffer containing rabbit anti-GFP antibody (1:500, A11122, Molecular Probes, USA) at 4 °C for 1–2 nights. After three 20 min washes with TBS-T, the sections were incubated with the secondary antibody donkey anti-rabbit IgG-AlexaFluor 488 (1:500, A21206, Molecular Probes, USA) in blocking buffer at room temperature for 3 h. Images were acquired using a Keyence microscope (BZ-9000, KEYENCE, Japan) with a Plan-Apochromat 4× or 10× objective lens.

### Synthetic data generation

We specified the time (T) of the data to be generated, the number of sequences to be embedded in the data (K), and the number of cells comprising one neuronal sequence (N_s_). Then, the number of cells in the synthetic data N was K × N_s_, and finally the original matrix V with size N × T was generated.

An exponential decay model was used to simulate the Ca^2+^ activity of neurons:

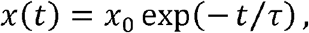

where x is the Ca^2+^ intensity of the cell, x_0_ is the initial Ca^2+^ intensity after the spike, and τ is the time constant. We expressed this in terms of a differential equation and updated it using the Euler method and also increased the Ca^2+^ intensity by 1 at each time point t with the firing probability p. To summarize the above, we obtained the updated equation:

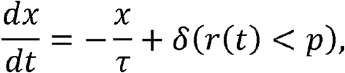

where δ is Kronecker’s delta function, and r is a random number following a uniform distribution *U* (0,1). We created K time series of x, with each randomly time-shifted from 0 to L, which is the time window of the neuronal sequences, to obtain the time series N_s_. By combining all of these, we created the original matrix V of synthetic data, with size N × T. The values of each parameter were: N_s_ = 100, τ = 2.0, p = 0.05, and L = 50.

### iSeq

#### Algorithm

iSeq is implemented in Python and operates on a GPU device. iSeq is a direct extension of convolutive NMF^19,20^. iSeq approximates the original matrix V of size N × T with dimensions of neurons and time, to the reconstructed matrix U with the same size and dimensions. U is computed by tensor convolution (see Math Note S1) of the pattern matrix W of size N × K × L with dimensions of neurons, number of neuronal sequences, the time window of neuronal sequences, and the intensity matrix H of size K × T+L−1 with dimensions of the number of neuronal sequences and time (Figures S3A–S3B). The number of neurons N and time T are determined from the shape of the original matrix V. The length of the time window of neuronal sequences L must be determined in advance. The number of neuronal sequences K is automatically determined by the method described in the subsection “Determination of K”.

The error between V and U is evaluated by the IS divergence^24^ and is expressed as follows:

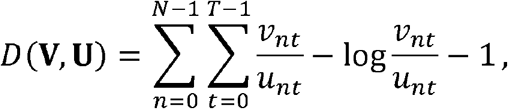

where ν*nt* and *unt* are values of V and U, respectively, at time *t* for the *n*^th^ neuron.

#### Multiplicative update rules

iSeq updates the values of the pattern matrix W and the intensity matrix H to decrease *D* (**V**,**U**). To guarantee the non-negativity of matrices, W and H are updated by the multiplicative update rule as follows^50^:

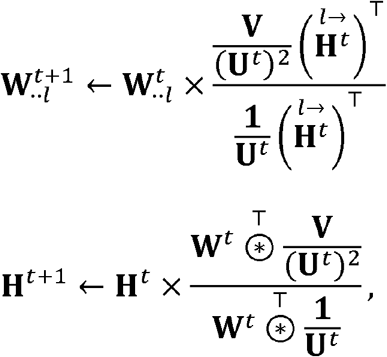

where ***W***^t^, H^t^, and ***U***^t^ are the *t*^*th*^ update results of the pattern matrix W, the intensity matrix H, and the reconstructed matrix U, respectively, and the symbol “×” and fraction represent the element-wise product and quotient of the matrix, respectively. For other symbols, see Math Note S1. However, if *D* (**V**,**U**) increases after the update, W and H are modified as follows:

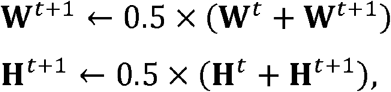

until *D* (**V**,**U**) decreases to guarantee convergence of W and H. See Math Note S2 for proof.

#### Determination of K

iSeq initially sets a sufficiently large value for K (=20). For the 20 detected neuronal sequences, it calculates the overlap matrix R, which is computed as transpose tensor convolution (Math Note S1) of the pattern matrix W and the original matrix V. Each row of R represents the time series of overlap (correlation) between each neuronal sequence and V. For each pair of rows in R, the Pearson’s correlation coefficient is calculated with a time shift of up to L−1 time frames both forward and backward, with the maximum value designated as the sequence similarity for that pair (see Figures 1D–1E). If the sequence similarity exceeds a predefined threshold, the pairs are considered to reflect the same neuronal sequences and are merged, resulting in a reduction K. This process is repeated until the sequence similarity between all remaining neuronal sequences falls below the threshold (see Figure 1F).

#### Determination of the threshold of similarity

We created 10 synthetic datasets, each with K, taking values from 1 to 10 and T = 3000. Ten datasets with five neuronal sequences (K=5) were decomposed by iSeq, varying the threshold of sequence similarity from 0.1 to 0.9. The decompositions with iSeq were performed 10 times for each dataset, using different random seeds. According to the number of neuronal sequences detected from the decomposition with the smallest reconstruction error *D* (**V**,**U**) in each calculation, five neuronal sequences were correctly detected when the correlation threshold was set within the range of 0.2–0.4. Therefore, we tentatively set the threshold of 0.3 and used the same method to decompose data containing 1–10 neuronal sequences. As a result, we decided to use 0.3 as the threshold because the correct number of neuronal sequences were detected in most of the datasets. Similar results were obtained for the datasets with T = 30000.

#### Matrix decomposition with compression

We set L = 50 time frames (10 s). In the beginning, the original matrix V was compressed by binning by L time frames (Figure S3C). We performed 30% of the entire iteration and decomposed this compressed matrix into the mini-pattern matrix W_mini_ of size N × K × 2 and the mini-intensity matrix H_mini_ of size K × T/L+1. The time frames for which the mini-intensity matrix H_mini_ had the largest value for each neuronal sequence were listed in order until they reached 10% of the total time frames of the compressed matrix (Figure S3D). The columns of the compressed matrix corresponding to these time frames were extracted and combined after unbinning them to create the high-density matrix (Figure S3E). Then, 30% of the entire iterations were performed, and this matrix was decomposed into the pattern matrix W of size N × K × L and the partial intensity matrix H_part_ of size K × T_part_, where T_part_ is the number of time frames of the high-density matrix (Figure S3G). The overlap matrix of the pattern matrix W and the high-density matrix was then calculated, and the appropriate value of K was determined based on the aforementioned method. Then, 10% of all iterations were performed to shape the remaining neuronal sequences. Finally, the original matrix V was restored by unbinning all columns of the compressed matrix (Figures S3F and S3H), and the remaining 30% of the total iterations were performed to obtain the final decomposition result (Figure S3I).

We performed the above calculations 10 times and used the decomposition result with the smallest reconstruction error *D* (**V**,**U**) in the subsequent analysis.

### Distance between trajectories

Let ***S*** = {***s***^1^, ***s***^*2*^,…, ***s***^m^} and ***F*** = {***f***^1^, ***f***^*2*^,…, ***f***^m^} represent the sets of trajectories corresponding to Success and Failure, respectively. Here, *m* and *n* are the number of trials that resulted in Success and Failure, respectively. 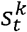 denotes the position of the mouse at time *t* during the *k*^th^ Success trial. The average Success trajectory is denoted by 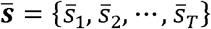, and the corresponding covariance matrices are denoted by ***∑*** = {***∑***_1_, ***∑***_*2*_,…, ***∑***_m_}, where

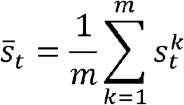

are satisfied, and *T* represents the time length of the trajectory. Similarly, the average Failure trajectory is denoted by 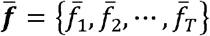, and the corresponding covariance matrices are denoted by ***Φ*** = {***Φ***_1_, ***Φ***_*2*_,…, ***Φ***_m_}. We calculated both the Euclidean distance

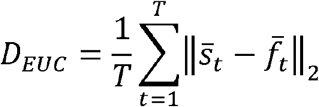

and Mahalanobis’ distance

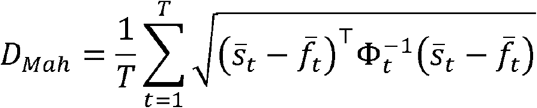

to evaluate the similarity between the trajectories ***S*** and ***F***. The Mahalanobis’ distance was compared using logarithmic transformations to normalize daily variances. Bartlett’s test was used without logarithmic transformation (\*\**P*<0.01, Figures 2G and 6F) and with logarithmic transformation (*P*=0.523, Figure 2G; *P*=0.374, Figure 6F).

### Sequence-sharpening operation

To eliminate redundancy and noise in the sequence representation in iSeq and convolutive NMF, the representation of the neuronal sequences in the pattern matrix W was modified to have no width of neural activity. Originally, a single neuronal sequence was represented by a two-dimensional matrix of size N × L (**W**_·k·_) within a three-dimensional pattern matrix W. However, we performed a manipulation such that for a single neuron, only one element in a row of length L would be positive, and the rest would be zero.

First, we reconstructed the neuronal activity produced by the k^th^ neuronal sequence U_k_ from **W**_·k·_ and part of the intensity matrix (**H**_k·_). Then, the sums of the time series (rows) of neuronal activity values were obtained based on U_k_. This value was the relative activity intensity w_int_ of the neurons composing the neuronal sequence, and the neuron with the largest value was referred to as the reference neuron. Subsequently, for the reference neuron and all other neurons, Pearson’s correlation was calculated for the activity time series represented in U_k_, with shifts of up to L−1 time frames both forward and backward. The amount of time shift that maximized the correlation coefficient was the relative activity timing w_tim_ of that neuron relative to the reference neuron within this neuronal sequence. We scaled the sum of w_int_ to be 1 and also adjusted w_tim_ to be within the length range of L. Thus, the shape of each neuronal sequence could now be represented by two vectors for each neuron: the activity intensity w_int_ and timing w_tim_. The values of the intensity matrix H were also adjusted to reconstruct approximately the same U, as the pattern matrix W was deformed.

### Spike intensity

We computed a moving average of 10 s before and after for the intensity of each neuronal sequence

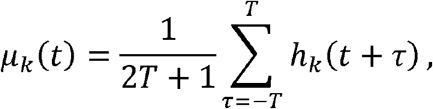

where *h*_*k*_(*t*) is the intensity of the *k*^*th*^ neuronal sequence at time *t* (corresponding to the elements of the *k*^*th*^ row and *t*^*th*^ column of the intensity matrix H), and *T* = 10 s. Based on this, we then scaled the intensity to compute the spike intensity of the neuronal sequence, as follows:

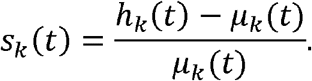

To avoid outliers, we modified the values of *S*_*k*_(*t*) to ensure they do not exceed 1.

### Location vector

We partitioned the Y-maze into 190 blocks, each 2 cm × 2 cm. The average spike intensity for the *k*^*th*^ neuronal sequences, when the mouse is in the *b*^*th*^ block can be calculated as follows:

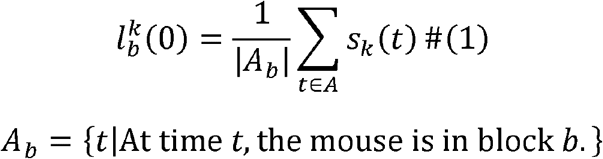

For the calculations, we applied a Gaussian filter covering up to 25 surrounding blocks to the mouse’s location and spike intensity data. The vector

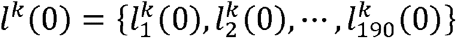

represents the posterior probability of spike intensity *s* given the mouse location *b, P(s\b)*. According to Bayes’ theorem

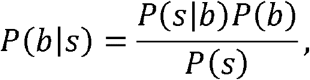

the likelihood of mouse location *b* given spike intensity *s* is proportional to this array.

Thus, using *l*^k^(0), we can estimate the distribution of the mouse’s location when the *k*^*th*^ neuronal sequence occurs.

We extended equation (1) to define the average spike intensity τ time frames before the mouse is located in the *b*^*th*^ block:

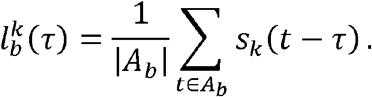

According to the discussion with Bayes’ theorem, we ascertain that vector

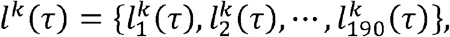

estimates the distribution of mouse location *b*, τ time frames after the *k*^th^ neuronal sequence occurs. Thus, the aggregated vector

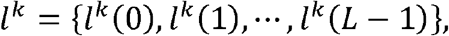

which combines all these vectors, predicts the mouse’s pathway during the activity period of the *k*^*th*^ neuronal sequence. We refer to this aggregated vector as the location vector.

### Decoding the mouse behaviors

#### Quantifying behaviors

Staying in the “Zone”: a value of 1 was assigned when the mouse was inside the Zone, and 0 was assigned when it was outside. Licking the “Port”: a value of 1 was assigned when the mouse licked the Port, and 0 was assigned when it did not. Acquiring a reward: a value of 1 was assigned when the mouse obtained a reward, otherwise 0 was assigned. The X- and Y-coordinates represent the mouse’s X- and Y-coordinates, respectively, on a scale from 0 to 1.

#### Decoding protocol

The decoder was implemented using the PyTorch library^51^ in Python, featuring an LSTM (long short-term memory) layer (with hidden_size=8, num_layers=2), a single fully connected layer, and a sigmoid function in the output layer. This decoder was trained to predict and output the behavior of the mouse at time *t*, based on the time series of the spike intensities of *K* neuronal sequences from *t* − 10.0 s to *t* − 0.2 s.

Decoding was conducted for each mouse (mouse IDs 1 to 6) and on specific days (days 1, 2, and 6). Separate decoders were set up and trained for each of the five behaviors described above.

For training, we used only the data from 25%–75% of the measured time (Training data), whereas the data from 0%–25% and 75%–100% of the time served as Validation data (Figure 7B). Throughout the training process, we continuously monitored prediction errors for both Training and Validation data. A typical sign of overfitting was observed when prediction errors on Training data decreased, whereas those on Validation data increased. Consequently, we selected the training results from the epoch in which the prediction error on Validation data was minimized as the final prediction error.

This training process was replicated 10 times for each behavior, and the overall prediction accuracy for each behavior was assessed based on the average of the final prediction errors.

#### Criteria of prediction (decoding) error

Let *Z* represent the set of behavioral data, and *Z*(*t*) denote the behavior at time *t*. As “staying in the ‘Zone’”, “licking the ‘Port’”, and “Acquiring reward” are binary behaviors, *Z*(*t*) takes values of 0 or 1. However, the frequency of *Z*(*t*) = 1 is considerably less than that of *Z*(*t*) = 0. To correct for this discrepancy, we quantify the error between the actual behavior *z* and the predicted behavior 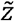 as follows:

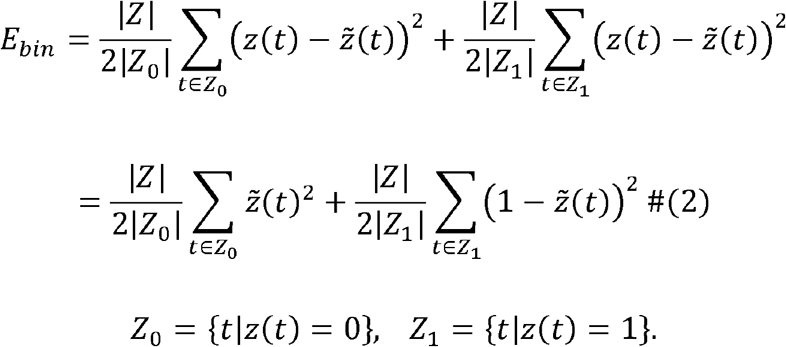

When the decoder consistently outputs a constant value *Z*(*t*) = *c*, this error function reaches its minimum value of 0.25|*Z*| at *c* = 0.5. We refer to the decoder that always outputs *Z*(*t*) =0.5 as the null decoder, and it serves as the baseline for assessing predictive performance.

By contrast, “X-coordinate” and “Y-coordinate” are continuous values ranging from 0 to

1. Here, we defined the error function for them as follows:

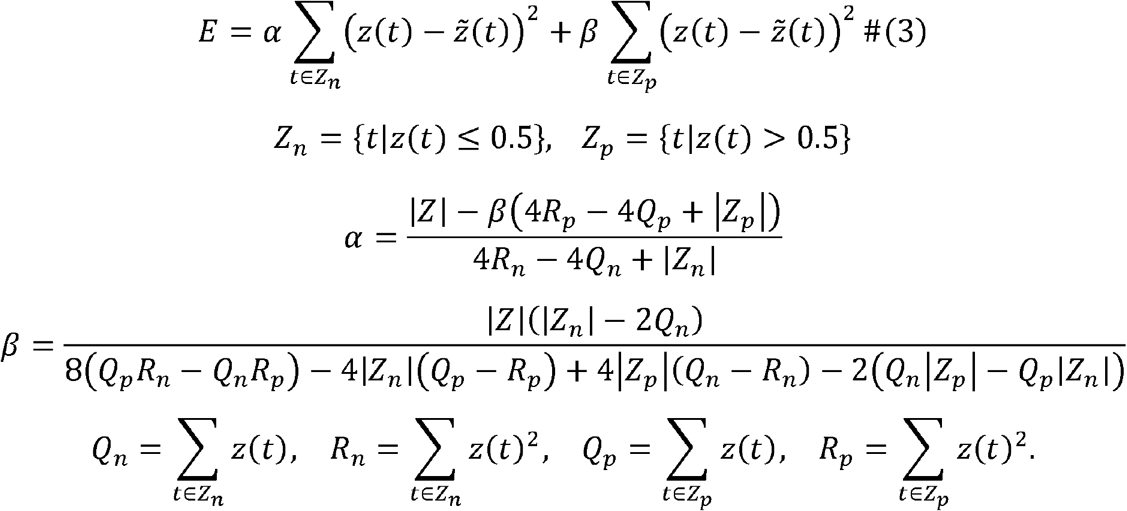

Remarkably, the error function for these coordinates also reaches a minimum value of 0.25|*Z*| when using the null decoder that always outputs *Z*(*t*) = 0.5. Considering *Z*(*t*) as a binary variable where *Z*_*n*_ = *Z*_*0*_, *Z*_*p*_ = *Z*_*1*_, *Q*_*n*_ = *R*_*0*_ = 0, *Q*_*p*_ = *R*_*p*_ = |*Z*_1_|, equation (3) simplifies to equation (2). This demonstrates that equation (3) is a natural extension of equation (2).

We employed the null decoder as a reference point and concluded that if the prediction error per data element significantly dropped below 0.25, this indicated that the neuronal sequences effectively captured behavioral information.

### Consistency of neuronal sequences

The consistency of neuronal sequences was assessed by the Pearson’s correlation between the similarity of cell populations and the similarity of pathways, as detailed below.

#### Similarity of cell populations

*W*^d^ was defined as the pattern matrix on day *d* after the sequence-sharpening operation, and *w*^d,k^ was defined as the relative activity intensity of the neurons consisting of *k*^*th*^ neuronal sequences. The values of *w*^d,k^ are scaled using its mean and standard deviation. The similarity between the cell populations of the *k*^*th*^ neuronal sequence on day *d* and the *κ*^*th*^ neuronal sequence on day δ is defined using cosine similarity as follows:

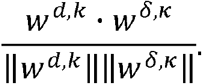

#### Similarity of pathways

We defined

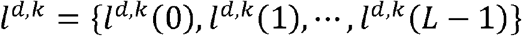

as the location vector of the *k*^*th*^ neuronal sequence on day *d*. The values of *l*^d,k^ are scaled using its mean and standard deviation. The similarity between the movement trajectory of the mouse during the activity of the *k*^*th*^ neuronal sequence on day *d* and the *κ*^*th*^ neuronal sequence on day δ was defined as the time-averaged cosine similarity:

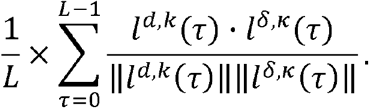

Additionally, considering that the location of the Zone was reversed on day 4, we also computed the pathway similarity for a transformed version of the location vector, where the block labels were adjusted by inverting the Y-coordinates.

