## Supplemental PDF for "The medial prefrontal cortex encodes procedural rules as sequential neuronal activity dynamics"

**Figure S1**

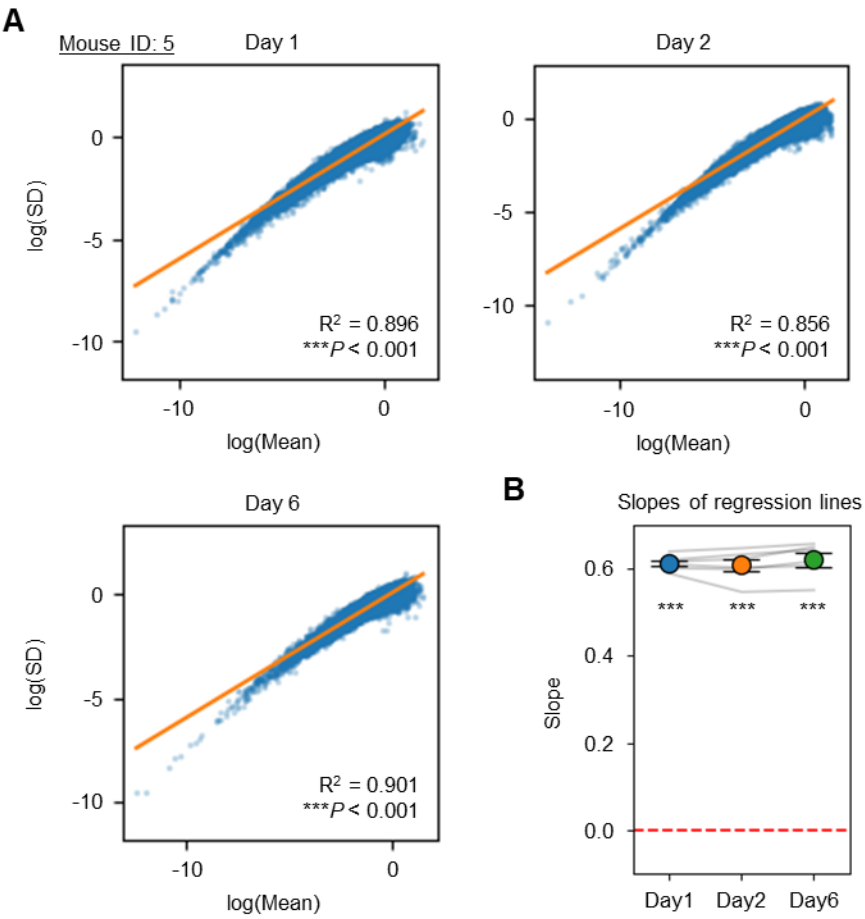


**Figure S1. $Ca^{2+}$ imaging data is influenced by signal-dependent noise, related to Figure 1.**

(A) Examples of the correlations between the mean values and standard deviations (SD) of the data. For each minute, we calculated the mean and SD of $Ca^{2+}$ signal intensities, then plotted these values on a logarithmic scale, as in Figure 1 of [S1]. Results for mouse ID: 5 on days 1, 2, and 6 are shown. The adjusted $R^2$ and P values are shown (Wald's test with t-distribution with Bonferroni's correction).

(B) Distribution of regression line slopes. Statistical tests were conducted to compare these slopes against zero (***$P$<0.001; one-sample t-test with Bonferroni's correction).

**Figure S2**

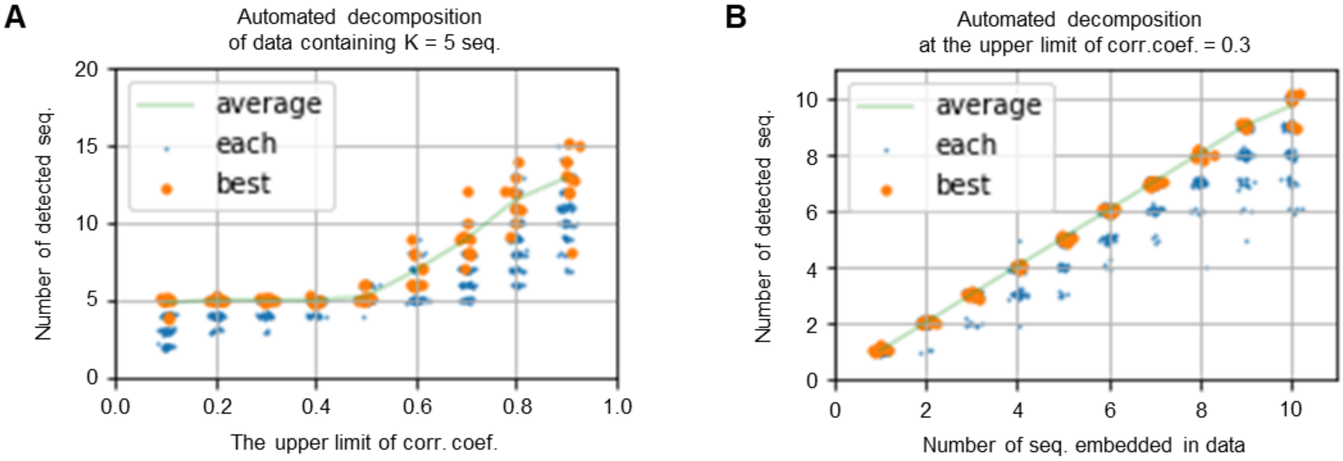


**Figure S2. Validation of decomposition accuracy of iSeq for synthetic data with T=30,000 time frames, related to Figure 1.**

(A, B) Results of decomposing synthetic data with T=30,000, conducted under the same conditions described in Figure 1H (A) and 1I (B).

**Figure S3**

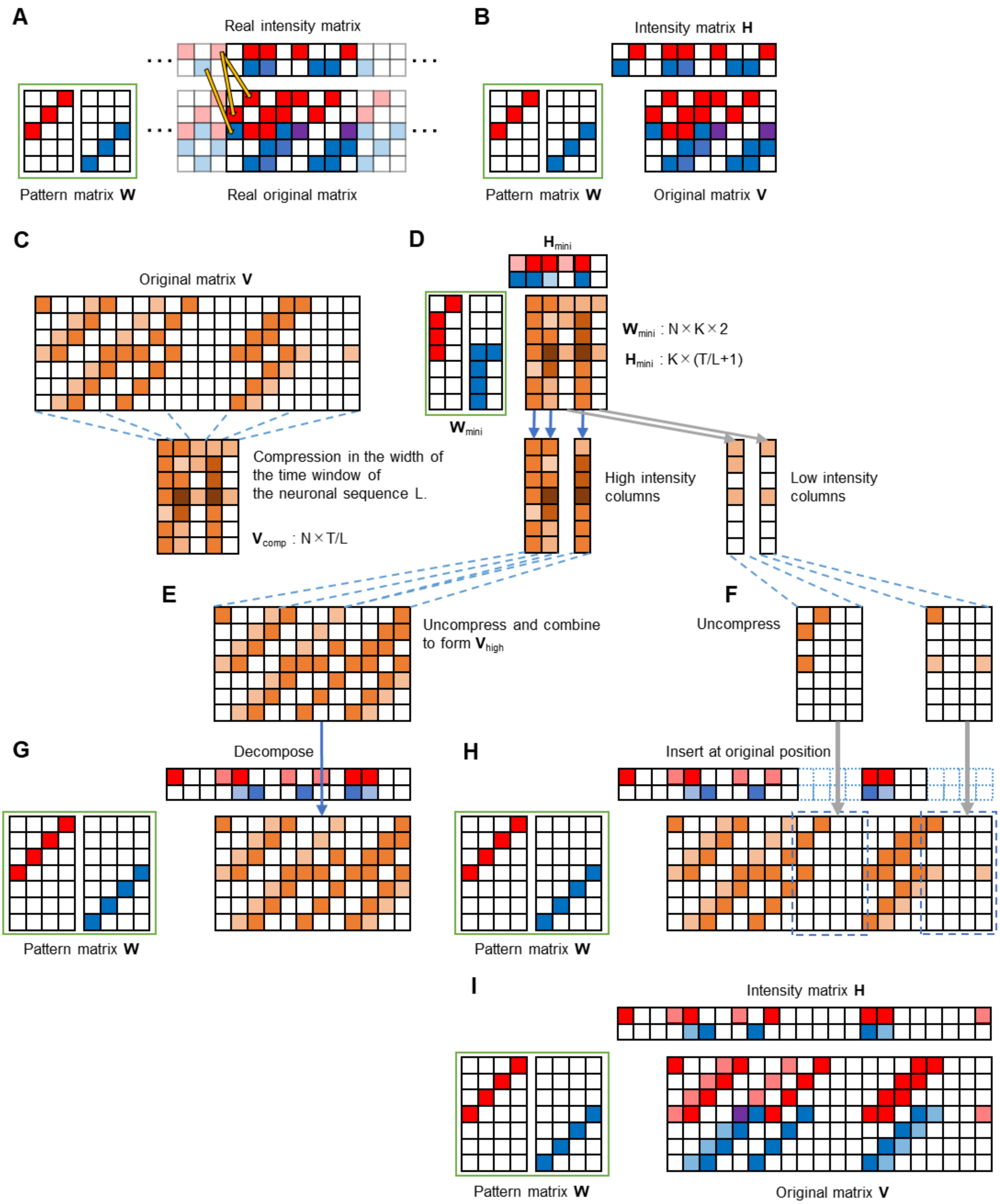


**Figure S3. Enhancements in the accuracy and acceleration of computations in the iSeq algorithm, related to Figure 1 and STAR Methods.**

(A) The initial portion of the original matrix V is influenced by neuronal sequence activities occurring outside the measured time frame.

(B) The shapes of the original matrix V, the pattern matrix W, and the intensity matrix H in the iSeq algorithm.

(C) Accelerating computations through matrix compression: the original matrix V is first compressed by binning according to the time window of neuronal sequence ($V_{comp}$).

(D) $V_{comp}$ is decomposed into matrices $W_{mini}$ and $H_{mini}$. The columns of $V_{comp}$ are sorted into high-intensity columns (here, top 60%) and low-intensity columns (here, bottom 40%) according to the values of $H_{mini}$.

(E) The high-intensity columns are decompressed and concatenated to form matrix $V_{high}$.

(F) The low-intensity columns are also decompressed.

(G) $V_{high}$ is decomposed to determine the pattern matrix W.

(H) The original matrix V is reconstructed by inserting the decompressed matrix from (F) into $V_{high}$.

(I) With W held constant, the intensity matrix H is recalculated to finalize the process.

**Figure S4**

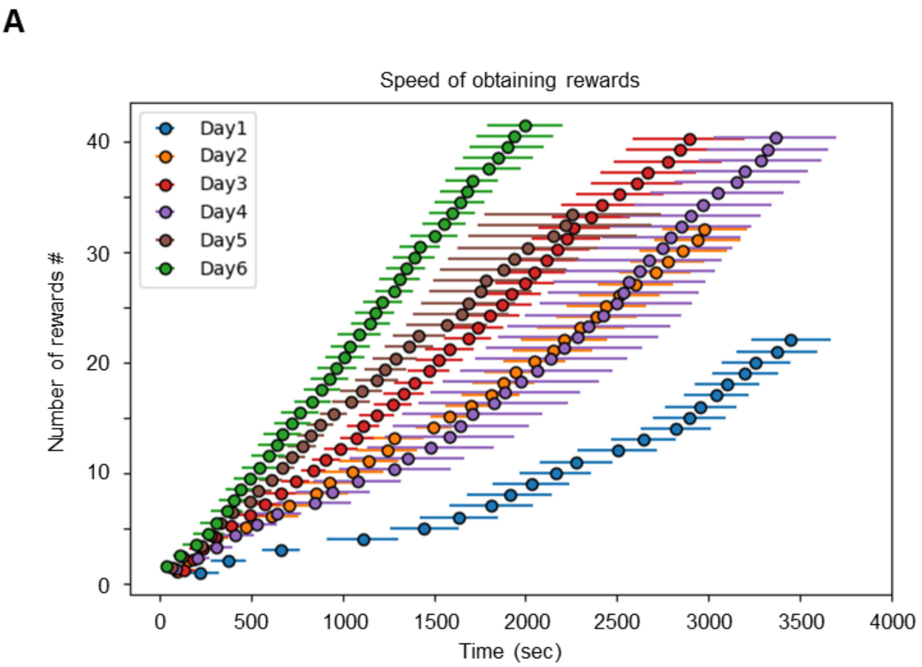


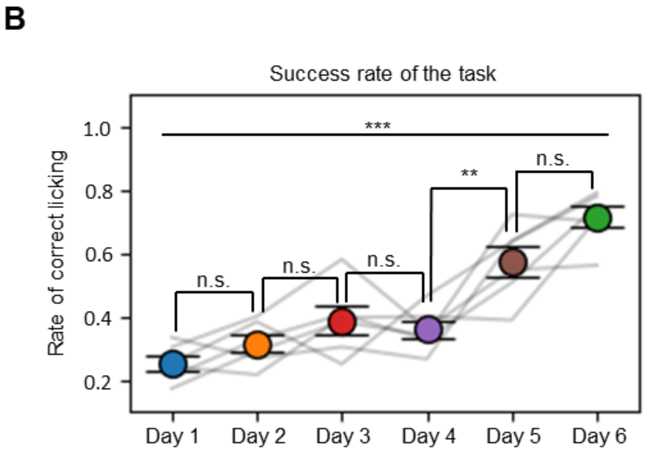


P-values

| | Day 1 | Day 2 | Day 3 | Day 4 | Day 5 | Day 6 |
|---|---|---|---|---|---|---|
| Day 1 | - | 0.8237 | 0.1188 | 0.3077 | <0.001 | <0.001 |
| Day 2 | | - | 0.7156 | 0.9451 | <0.001 | <0.001 |
| Day 3 | | | - | 0.9947 | 0.0111 | <0.001 |
| Day 4 | | | | - | 0.0028 | <0.001 |
| Day 5 | | | | | - | 0.0902 |
| Day 6 | | | | | | - |

**Figure S4. Task performance of mice during the entire training period, related to Figure 2.**

(A) Same as Figure 2C, but for days 1–6.

(B) Same as Figure 2D, but for days 1–6 (left). Post hoc P values (right) for all combinations of pairs (one-way RM ANOVA with Tukey's post hoc test).

**Figure S5**

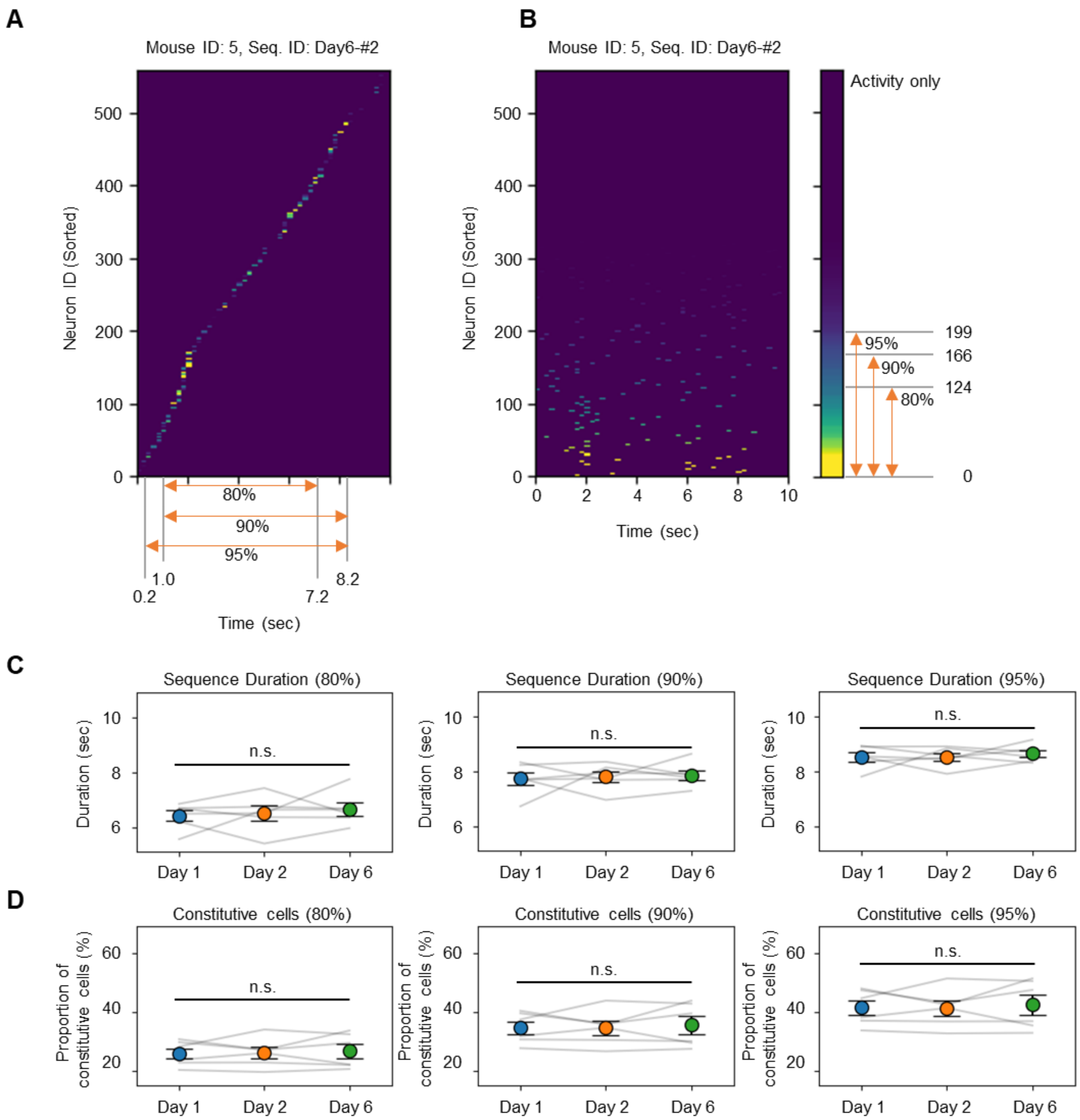


**Figure S5. Definition of the duration and number of constitutive cells of neuronal sequences, related to Figure 3.**

(A) Definition of duration. Neurons were sorted by the order of their activity timing within the neuronal sequence. Then the duration of the neuronal sequence was defined as the shortest period during which the sum of activity values reached 80%, 90%, and 95% of the total activity.

(B) Definition of the number of constitutive cells. Neurons were sorted by the order of their activity value within the neuronal sequence. Then the number of constitutive cells was defined as the smallest number of cells for which the sum of activity values reached 80%, 90%, and 95% of the total activity.

(C) Duration of detected neuronal sequences for each day of the experiment (80%, $P$=0.73; 90%, $P$=0.90; 95%, $P$=0.79; one-way RM ANOVA).

(D) Same as (C), but for the proportion of constitutive cells relative to all cells in the measured field of $Ca^{2+}$ imaging

(80%, *P*=0.81; 90%, *P*=0.79; 95%, *P*=0.80; one-way RM ANOVA).

These calculations were performed after the sequence-sharpening operation (see Figures S5D–S5E).

Figure S6

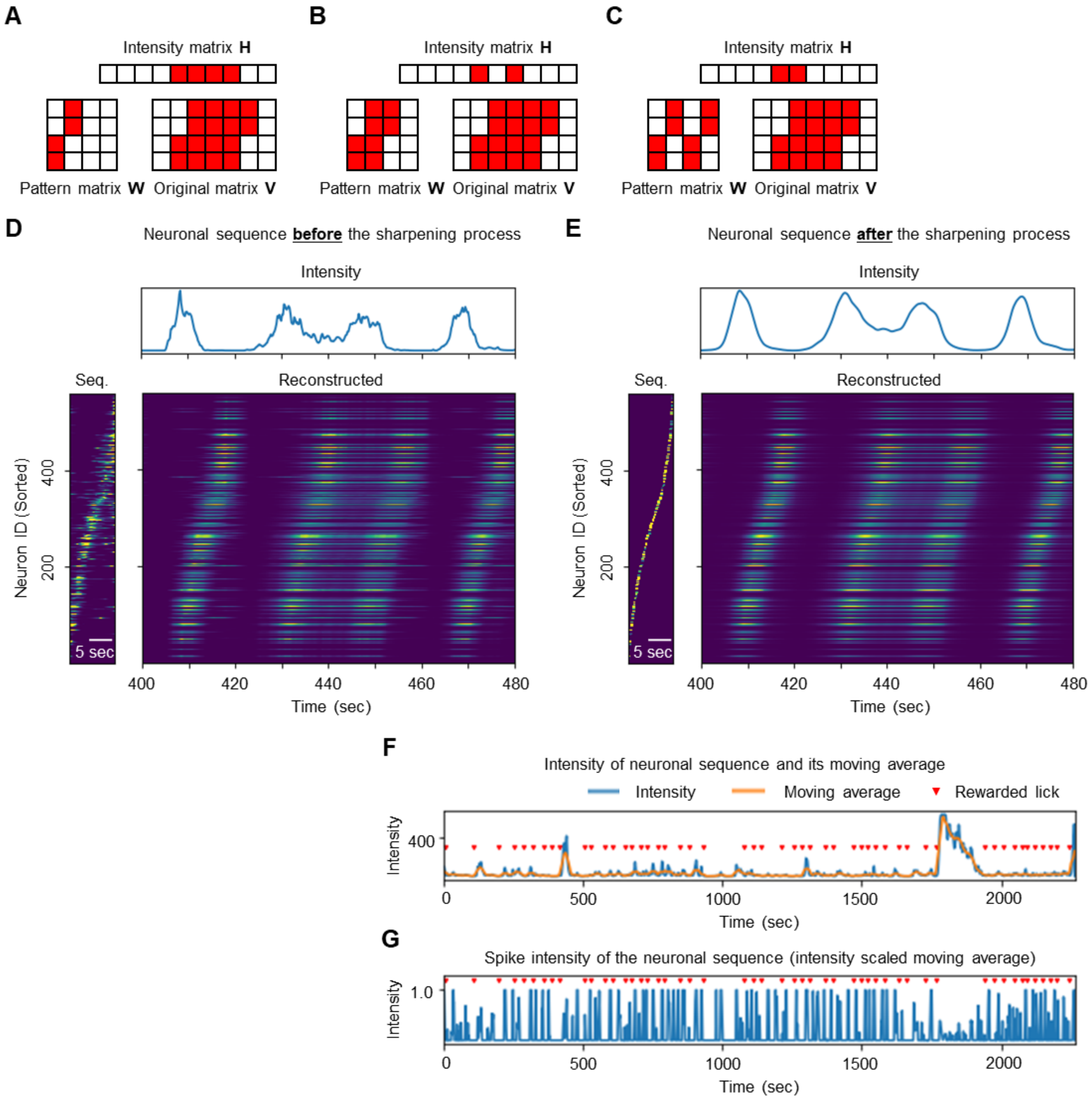


**Figure S6. The sequence-sharpening operation and spike intensity, related to Figure 4.**

(A–C) An example of the same original matrix V being decomposed into (A) “narrow”, (B) “wider”, and (C) “separated” sequences.

(D–E) An example of before (D) and after (E) the sequence-sharpening operation. The following is shown: the intensity of the neuronal sequence (top), the shape of the neuronal sequence (bottom left), and the neuronal activity reconstructed from the shape and intensity of the neuronal sequence (bottom right).

(F) An example of the intensity of a neuronal sequence and its moving average.

(G) The spike intensity calculated from (F).

**Figures S7**

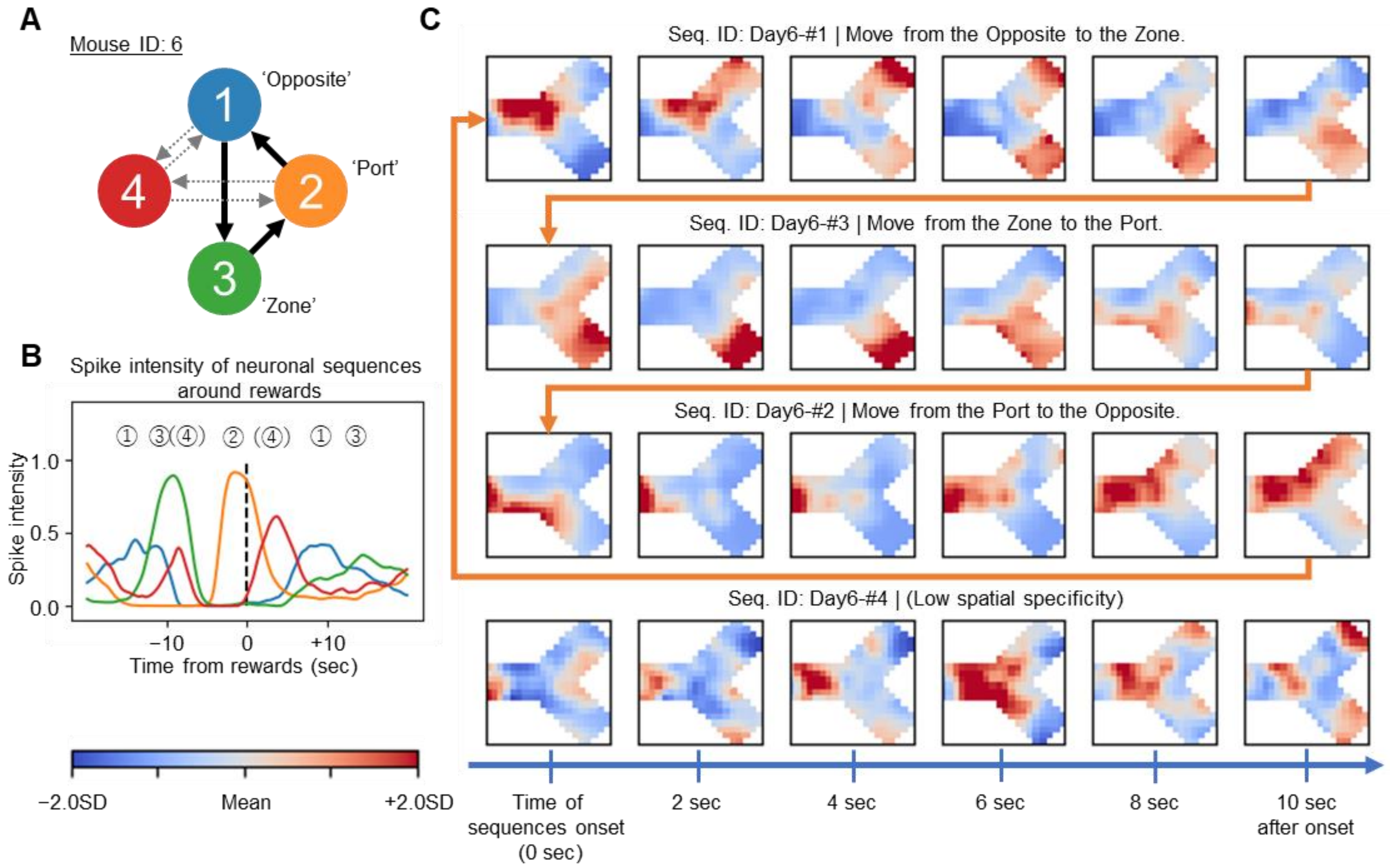


**Figures S7. Neuronal sequences of the mouse with a three-step action comprehension of the rule, related to Figure 4.**

(A) The relations between neuronal sequences and rule components of mouse ID: 6.

(B) Time series of mean spike intensities of neuronal sequences before and after obtaining a reward. The number at the top represents the corresponding sequence ID.

(C) Same as Figure 4D, but for mouse ID: 6.

| Mouse ID | Seq. ID | | Duration (sec) | | | Seq. ID | | Duration (sec) | | | Seq. ID | | Duration (sec) | | |
|---|---|---|---|---|---|---|---|---|---|---|---|---|---|---|---|
| | | | 80% | 90% | 95% | | | 80% | 90% | 95% | | | 80% | 90% | 95% |
| 1 | Day1 | #1 | 3.4 | 8.4 | 9.2 | Day2 | #1 | 3.8 | 6.4 | 7.8 | Day6 | #1 | 6.6 | 7.6 | 9.0 |
| | | #2 | 9.0 | 9.4 | 9.4 | | #2 | 5.4 | 7.0 | 8.0 | | #2 | 5.2 | 6.8 | 7.2 |
| | | #3 | 7.2 | 8.0 | 9.0 | | #3 | 5.2 | 6.2 | 7.2 | | #3 | 6.6 | 8.0 | 8.6 |
| | | #4 | 4.8 | 5.8 | 7.4 | | #4 | 7.2 | 8.2 | 8.6 | | #4 | 5.2 | 6.0 | 8.2 |
| | | #5 | 7.0 | 8.8 | 9.4 | | | | | | | #5 | 6.2 | 8.0 | 8.8 |
| | | #6 | 5.8 | 6.0 | 6.8 | | | | | | | | | | |
| | Average | | 6.2 | 7.7 | 8.5 | Average | | 5.4 | 7.0 | 7.9 | Average | | 6.0 | 7.3 | 8.4 |
| | | | | | | | | | | | Ave. Mouse ID: 1 | | 5.9 | 7.4 | 8.3 |

| Mouse ID | Seq. ID | | Duration (sec) | | | Seq. ID | | Duration (sec) | | | Seq. ID | | Duration (sec) | | |
|---|---|---|---|---|---|---|---|---|---|---|---|---|---|---|---|
| | | | 80% | 90% | 95% | | | 80% | 90% | 95% | | | 80% | 90% | 95% |
| 2 | Day1 | #1 | 7.4 | 9.2 | 9.4 | Day2 | #1 | 7.8 | 8.2 | 8.6 | Day6 | #1 | 6.2 | 7.2 | 8.2 |
| | | #2 | 7.6 | 8.8 | 9.2 | | #2 | 7.8 | 8.8 | 9.0 | | #2 | 7.4 | 8.8 | 9.2 |
| | | #3 | 7.0 | 8.2 | 8.8 | | #3 | 7.0 | 8.2 | 8.8 | | #3 | 4.8 | 7.6 | 8.6 |
| | | #4 | 6.6 | 8.6 | 9.0 | | #4 | 4.6 | 5.8 | 7.8 | | #4 | 6.2 | 7.6 | 8.6 |
| | | #5 | 6.8 | 7.6 | 9.2 | | #5 | 8.6 | 9.2 | 9.4 | | #5 | 7.0 | 8.6 | 9.2 |
| | | #6 | 7.4 | 8.4 | 8.8 | | #6 | 7.6 | 8.8 | 9.0 | | #6 | 7.2 | 8.4 | 8.8 |
| | | #7 | 5.8 | 7.4 | 8.6 | | #7 | 8.0 | 8.8 | 9.2 | | #7 | 6.2 | 6.4 | 8.0 |
| | | #8 | 7.8 | 8.8 | 9.0 | | #8 | 6.6 | 7.6 | 8.6 | | #8 | 6.8 | 8.2 | 9.0 |
| | | #9 | 5.2 | 7.0 | 8.0 | | #9 | 7.8 | 8.8 | 9.2 | | | | | |
| | | | | | | | #10 | 8.4 | 9.2 | 9.4 | | | | | |
| | Average | | 6.8 | 8.2 | 8.9 | Average | | 7.4 | 8.3 | 8.9 | Average | | 6.5 | 7.8 | 8.7 |
| | | | | | | | | | | | Avg. Mouse ID: 2 | | 6.9 | 8.2 | 8.8 |

| Mouse ID | Seq. ID | | Duration (sec) | | | Seq. ID | | Duration (sec) | | | Seq. ID | | Duration (sec) | | |
|---|---|---|---|---|---|---|---|---|---|---|---|---|---|---|---|
| | | | 80% | 90% | 95% | | | 80% | 90% | 95% | | | 80% | 90% | 95% |
| 3 | Day1 | #1 | 5.4 | 6.4 | 7.2 | Day2 | #1 | 5.6 | 7.0 | 7.4 | Day6 | #1 | 7.4 | 8.8 | 9.2 |
| | | #2 | 5.4 | 7.0 | 8.0 | | #2 | 6.0 | 6.8 | 7.0 | | #2 | 7.2 | 8.0 | 8.8 |
| | | #3 | 7.6 | 8.4 | 9.0 | | #3 | 6.2 | 7.6 | 8.6 | | #3 | 5.8 | 7.8 | 8.8 |
| | | #4 | 6.8 | 8.0 | 8.6 | | #4 | 8.4 | 9.0 | 9.4 | | #4 | 7.0 | 7.8 | 8.4 |
| | | #5 | 6.6 | 7.4 | 7.8 | | #5 | 6.8 | 8.8 | 9.2 | | #5 | 5.8 | 7.2 | 8.8 |
| | | #6 | 7.2 | 8.0 | 8.8 | | #6 | 7.6 | 8.4 | 8.8 | | #6 | 7.2 | 8.2 | 8.6 |
| | | #7 | 7.8 | 8.6 | 9.0 | | #7 | 6.0 | 8.0 | 9.0 | | #7 | 6.4 | 7.4 | 8.6 |
| | | | | | | | | | | | | #8 | 6.6 | 8.0 | 9.2 |
| | Average | | 6.7 | 7.7 | 8.3 | Average | | 6.7 | 7.9 | 8.5 | Average | | 6.7 | 7.9 | 8.8 |
| | | | | | | | | | | | Avg. Mouse ID: 3 | | 6.7 | 7.8 | 8.6 |

| Mouse ID | Seq. ID | | Duration (sec) | | | Seq. ID | | Duration (sec) | | | Seq. ID | | Duration (sec) | | |
|---|---|---|---|---|---|---|---|---|---|---|---|---|---|---|---|
| | | | 80% | 90% | 95% | | | 80% | 90% | 95% | | | 80% | 90% | 95% |
| 4 | Day1 | #1 | 7.6 | 8.4 | 8.8 | Day2 | #1 | 5.0 | 6.4 | 7.4 | Day6 | #1 | 8.2 | 9.4 | 9.6 |
| | | #2 | 6.0 | 7.6 | 8.4 | | #2 | 6.2 | 8.0 | 8.8 | | #2 | 8.2 | 8.8 | 9.2 |
| | | #3 | 7.6 | 8.4 | 8.8 | | #3 | 6.8 | 8.2 | 8.8 | | #3 | 8.8 | 9.4 | 9.6 |
| | | #4 | 7.0 | 8.0 | 8.6 | | #4 | 8.0 | 8.8 | 9.2 | | #4 | 8.2 | 8.8 | 9.2 |
| | | #5 | 5.6 | 7.6 | 8.0 | | #5 | 6.4 | 7.6 | 8.2 | | #5 | 7.4 | 8.4 | 8.8 |
| | | #6 | 5.8 | 7.2 | 8.4 | | #6 | 6.6 | 7.4 | 8.2 | | #6 | 7.8 | 8.4 | 8.8 |
| | | #7 | 5.8 | 6.6 | 8.8 | | | | | | | #7 | 5.6 | 7.2 | 8.8 |
| | Average | | 6.5 | 7.7 | 8.5 | Average | | 6.5 | 7.7 | 8.4 | Average | | 7.7 | 8.6 | 9.1 |
| | | | | | | | | | | | Avg. Mouse ID: 4 | | 6.9 | 8.0 | 8.7 |

| Mouse ID | Seq. ID | | Duration (sec) | | | Seq. ID | | Duration (sec) | | | Seq. ID | | Duration (sec) | | |
|---|---|---|---|---|---|---|---|---|---|---|---|---|---|---|---|
| | | | 80% | 90% | 95% | | | 80% | 90% | 95% | | | 80% | 90% | 95% |
| 5 | Day1 | #1 | 5.0 | 6.2 | 7.0 | Day2 | #1 | 5.8 | 7.2 | 8.0 | Day6 | #1 | 8.0 | 8.8 | 9.2 |
| | | #2 | 6.8 | 7.6 | 8.4 | | #2 | 6.8 | 8.2 | 8.4 | | #2 | 6.2 | 7.2 | 8.0 |
| | | #3 | 6.8 | 8.0 | 8.4 | | #3 | 4.2 | 7.2 | 9.2 | | #3 | 4.6 | 6.4 | 7.4 |
| | | #4 | 3.8 | 5.0 | 6.6 | | #4 | 6.4 | 8.0 | 8.8 | | #4 | 8.0 | 8.8 | 9.4 |
| | | #5 | 5.4 | 6.8 | 8.6 | | #5 | 8.6 | 9.2 | 9.4 | | #5 | 6.4 | 7.6 | 8.6 |
| | | | | | | | #6 | 8.0 | 9.0 | 9.2 | | | | | |
| | Average | | 5.6 | 6.7 | 7.8 | Average | | 6.6 | 8.1 | 8.8 | Average | | 6.6 | 7.8 | 8.5 |
| | | | | | | | | | | | Avg. Mouse ID: 5 | | 6.3 | 7.6 | 8.4 |

| Mouse ID | Seq. ID | | Duration (sec) | | | Seq. ID | | Duration (sec) | | | Seq. ID | | Duration (sec) | | |
|---|---|---|---|---|---|---|---|---|---|---|---|---|---|---|---|
| | | | 80% | 90% | 95% | | | 80% | 90% | 95% | | | 80% | 90% | 95% |
| 6 | Day1 | #1 | 7.4 | 8.2 | 8.4 | Day2 | #1 | 7.8 | 8.8 | 9.2 | Day6 | #1 | 7.6 | 8.4 | 8.6 |
| | | #2 | 6.6 | 7.8 | 8.8 | | #2 | 5.4 | 7.0 | 8.4 | | #2 | 6.0 | 7.0 | 7.6 |
| | | #3 | 6.4 | 8.2 | 8.8 | | #3 | 6.6 | 7.6 | 8.6 | | #3 | 6.4 | 7.4 | 8.0 |
| | | #4 | 6.0 | 8.8 | 9.4 | | #4 | 4.8 | 7.0 | 8.2 | | #4 | 5.4 | 8.0 | 9.0 |
| | | #5 | 7.0 | 8.6 | 9.2 | | #5 | 7.2 | 8.0 | 8.4 | | | | | |
| | Average | | 6.7 | 8.3 | 8.9 | Average | | 6.4 | 7.7 | 8.6 | Average | | 6.3 | 7.7 | 8.3 |
| | | | | | | | | | | | Avg. Mouse ID: 6 | | 6.5 | 7.9 | 8.6 |

| | Duration (sec) | | | | Duration (sec) | | | | Duration (sec) | | |
|---|---|---|---|---|---|---|---|---|---|---|---|
| | 80% | 90% | 95% | | 80% | 90% | 95% | | 80% | 90% | 95% |
| Avg. Day1 | 6.5 | 7.8 | 8.5 | Avg. Day2 | 6.7 | 7.9 | 8.6 | Avg. Day6 | 6.7 | 7.9 | 8.7 |
| | | | | | | | | Avg. Total | 6.6 | 7.9 | 8.6 |

**Table S1. The duration of neuronal sequences, related to Figure 3.**

| Mouse ID | Seq. ID | | Number of Cells | | | Seq. ID | | Number of Cells | | | Seq. ID | | Number of Cells | | |
|---|---|---|---|---|---|---|---|---|---|---|---|---|---|---|---|
| | | | 80% | 90% | 95% | | | 80% | 90% | 95% | | | 80% | 90% | 95% |
| 1 | Day1 | #1 | 98 | 126 | 149 | Day2 | #1 | 122 | 157 | 182 | Day6 | #1 | 89 | 123 | 148 |
| | | #2 | 105 | 140 | 168 | | #2 | 112 | 146 | 172 | | #2 | 101 | 143 | 174 |
| Total Number of Cells | | #3 | 86 | 117 | 143 | | #3 | 128 | 161 | 186 | | #3 | 131 | 168 | 193 |
| | | #4 | 118 | 154 | 180 | | #4 | 110 | 144 | 172 | | #4 | 111 | 144 | 171 |
| | | #5 | 105 | 140 | 165 | | | | | | | #5 | 130 | 164 | 189 |
| 347 | | #6 | 76 | 102 | 124 | | | | | | | | | | |
| | Average | | 98.0 | 129.8 | 154.8 | Average | | 118.0 | 152.0 | 178.0 | Average | | 112.4 | 148.4 | 175.0 |
| | | | | | | | | | | | Ave. Mouse ID: 1 | | 108.1 | 141.9 | 167.7 |

| Mouse ID | Seq. ID | | Number of Cells | | | Seq. ID | | Number of Cells | | | Seq. ID | | Number of Cells | | |
|---|---|---|---|---|---|---|---|---|---|---|---|---|---|---|---|
| | | | 80% | 90% | 95% | | | 80% | 90% | 95% | | | 80% | 90% | 95% |
| 2 | Day1 | #1 | 71 | 100 | 126 | Day2 | #1 | 133 | 176 | 212 | Day6 | #1 | 124 | 159 | 184 |
| | | #2 | 98 | 135 | 165 | | #2 | 93 | 126 | 154 | | #2 | 89 | 122 | 150 |
| Total Number of Cells | | #3 | 116 | 154 | 181 | | #3 | 89 | 119 | 146 | | #3 | 110 | 146 | 174 |
| | | #4 | 123 | 165 | 199 | | #4 | 52 | 74 | 98 | | #4 | 95 | 126 | 149 |
| | | #5 | 70 | 106 | 138 | | #5 | 93 | 128 | 156 | | #5 | 93 | 124 | 148 |
| 455 | | #6 | 89 | 120 | 146 | | #6 | 92 | 123 | 148 | | #6 | 67 | 92 | 111 |
| | | #7 | 94 | 125 | 152 | | #7 | 87 | 121 | 152 | | #7 | 87 | 116 | 141 |
| | | #8 | 82 | 109 | 131 | | #8 | 91 | 120 | 146 | | #8 | 85 | 114 | 139 |
| | | #9 | 88 | 119 | 142 | | #9 | 83 | 117 | 144 | | | | | |
| | | | | | | | #10 | 78 | 107 | 131 | | | | | |
| | Average | | 92.3 | 125.9 | 153.3 | Average | | 89.1 | 121.1 | 148.7 | Average | | 93.8 | 124.9 | 149.5 |
| | | | | | | | | | | | Avg. Mouse ID: 2 | | 91.6 | 123.8 | 150.5 |

| Mouse ID | Seq. ID | | Number of Cells | | | Seq. ID | | Number of Cells | | | Seq. ID | | Number of Cells | | |
|---|---|---|---|---|---|---|---|---|---|---|---|---|---|---|---|
| | | | 80% | 90% | 95% | | | 80% | 90% | 95% | | | 80% | 90% | 95% |
| 3 | Day1 | #1 | 115 | 158 | 193 | Day2 | #1 | 168 | 226 | 272 | Day6 | #1 | 110 | 149 | 182 |
| | | #2 | 151 | 194 | 226 | | #2 | 148 | 197 | 235 | | #2 | 91 | 122 | 153 |
| Total Number of Cells | | #3 | 117 | 161 | 198 | | #3 | 132 | 172 | 210 | | #3 | 154 | 211 | 255 |
| | | #4 | 140 | 192 | 232 | | #4 | 109 | 144 | 173 | | #4 | 179 | 229 | 264 |
| | | #5 | 124 | 162 | 193 | | #5 | 113 | 149 | 178 | | #5 | 128 | 182 | 225 |
| 546 | | #6 | 110 | 152 | 189 | | #6 | 96 | 125 | 148 | | #6 | 101 | 149 | 191 |
| | | #7 | 113 | 152 | 183 | | #7 | 106 | 151 | 188 | | #7 | 103 | 143 | 179 |
| | | | | | | | | | | | | #8 | 94 | 128 | 159 |
| | Average | | 124.3 | 167.3 | 202.0 | Average | | 124.6 | 166.3 | 200.6 | Average | | 120.0 | 164.1 | 201.0 |
| | | | | | | | | | | | Avg. Mouse ID: 3 | | 122.8 | 165.8 | 201.2 |

| Mouse ID | Seq. ID | | Number of Cells | | | Seq. ID | | Number of Cells | | | Seq. ID | | Number of Cells | | |
|---|---|---|---|---|---|---|---|---|---|---|---|---|---|---|---|
| | | | 80% | 90% | 95% | | | 80% | 90% | 95% | | | 80% | 90% | 95% |
| 4 | Day1 | #1 | 115 | 160 | 195 | Day2 | #1 | 181 | 243 | 290 | Day6 | #1 | 103 | 136 | 165 |
| | | #2 | 142 | 184 | 218 | | #2 | 118 | 162 | 195 | | #2 | 118 | 158 | 188 |
| Total Number of Cells | | #3 | 155 | 205 | 246 | | #3 | 147 | 189 | 221 | | #3 | 145 | 189 | 223 |
| | | #4 | 127 | 167 | 199 | | #4 | 185 | 244 | 288 | | #4 | 116 | 155 | 188 |
| | | #5 | 151 | 208 | 254 | | #5 | 142 | 188 | 226 | | #5 | 117 | 159 | 194 |
| 568 | | #6 | 103 | 137 | 165 | | #6 | 114 | 154 | 187 | | #6 | 171 | 227 | 271 |
| | | #7 | 150 | 202 | 239 | | | | | | | #7 | 111 | 149 | 178 |
| | Average | | 134.7 | 180.4 | 216.6 | Average | | 147.8 | 196.7 | 234.5 | Average | | 125.9 | 167.6 | 201.0 |
| | | | | | | | | | | | Avg. Mouse ID: 4 | | 135.6 | 180.8 | 216.5 |

| Mouse ID | Seq. ID | | Number of Cells | | | Seq. ID | | Number of Cells | | | Seq. ID | | Number of Cells | | |
|---|---|---|---|---|---|---|---|---|---|---|---|---|---|---|---|
| | | | 80% | 90% | 95% | | | 80% | 90% | 95% | | | 80% | 90% | 95% |
| 5 | Day1 | #1 | 162 | 217 | 258 | Day2 | #1 | 194 | 253 | 300 | Day6 | #1 | 166 | 224 | 270 |
| | | #2 | 186 | 246 | 295 | | #2 | 171 | 227 | 271 | | #2 | 124 | 166 | 199 |
| Total Number of Cells | | #3 | 127 | 173 | 212 | | #3 | 146 | 190 | 224 | | #3 | 160 | 224 | 278 |
| | | #4 | 200 | 258 | 302 | | #4 | 122 | 165 | 201 | | #4 | 193 | 253 | 301 |
| | | #5 | 155 | 210 | 255 | | #5 | 156 | 205 | 243 | | #5 | 181 | 236 | 277 |
| 559 | | | | | | | #6 | 121 | 166 | 207 | | | | | |
| | Average | | 166.0 | 220.8 | 264.4 | Average | | 151.7 | 201.0 | 241.0 | Average | | 164.8 | 220.6 | 265.0 |
| | | | | | | | | | | | Avg. Mouse ID: 5 | | 160.2 | 213.3 | 255.8 |

| Mouse ID | Seq. ID | | Number of Cells | | | Seq. ID | | Number of Cells | | | Seq. ID | | Number of Cells | | |
|---|---|---|---|---|---|---|---|---|---|---|---|---|---|---|---|
| | | | 80% | 90% | 95% | | | 80% | 90% | 95% | | | 80% | 90% | 95% |
| 6 | Day1 | #1 | 265 | 349 | 415 | Day2 | #1 | 171 | 230 | 278 | Day6 | #1 | 263 | 341 | 400 |
| | | #2 | 183 | 245 | 295 | | #2 | 197 | 262 | 313 | | #2 | 259 | 336 | 392 |
| Total Number of Cells | | #3 | 235 | 306 | 361 | | #3 | 175 | 241 | 290 | | #3 | 260 | 336 | 393 |
| | | #4 | 216 | 283 | 335 | | #4 | 223 | 282 | 327 | | #4 | 199 | 263 | 311 |
| | | #5 | 220 | 287 | 340 | | #5 | 221 | 287 | 337 | | | | | |
| 729 | Average | | 223.8 | 294.0 | 349.2 | Average | | 197.4 | 260.4 | 309.0 | Average | | 245.2 | 319.0 | 374.0 |
| | | | | | | | | | | | Avg. Mouse ID: 6 | | 220.5 | 289.1 | 341.9 |

| | Number of Cells | | | | Number of Cells | | | | Number of Cells | | |
|---|---|---|---|---|---|---|---|---|---|---|---|
| | 80% | 90% | 95% | | 80% | 90% | 95% | | 80% | 90% | 95% |
| Avg. Day1 | 132.8 | 177.4 | 213.0 | Avg. Day2 | 132.1 | 175.6 | 210.6 | Avg. Day6 | 134.0 | 178.5 | 213.7 |
| | | | | | | | | Avg. Total | 133.0 | 177.2 | 212.4 |

**Table S2. The number of constitutive cells of neuronal sequences, related to Figure 3.**

**Math Note S1. Definitions of operators used in this paper, related to Figure 1.**

According to Mackevicius et al. 2019, we defined each operator as follows.

**Matrix indexing.** Any index of a vector with length T is specified by t. This case relationship applies to all letters. Additionally, when indexing specifically extracts elements of a matrix, the elements are represented by the lower-case letter corresponding to the letter representing the matrix. In other words, the element in row n and column t of the original matrix V is $v_{nt}$.

To accommodate negative indexing, which is discussed below, 0-based indexing is applied throughout this article. This means that vectors and rows and columns of matrices of length T are assigned an index number from 0 to T−1. In addition, we use · to represent all elements along each dimension of the matrix. Therefore, $\mathbf{V}_{\cdot t}$ means the vector of $\{v_{0,t}, v_{1,t}, \cdots, v_{N-1,t}\}$. For the pattern matrix W, which is a 3D matrix, $\mathbf{W}_{\cdot k \cdot}$ is a 2D matrix and represents the shape of the k^th^ neuronal sequence.

The intensity matrix H is partially negatively indexed because it is extended forward in the time dimension by L−1. This means that the range of indices in the time dimension of H is −L+1 to T−1.

**Shift operator.** The operator $\overset{l\rightarrow}{\mathbf{V}}$ shifts $\mathbf{V}$ in the direction of → by $l$ time frames. In other words, the correspondence is as follows:

$$\left(\overset{l\rightarrow}{\mathbf{V}}\right)_{\cdot t} = \mathbf{V}_{\cdot (t-l)}$$

$$\left(\overset{\leftarrow l}{\mathbf{V}}\right)_{\cdot t} = \mathbf{V}_{\cdot (t+l)}$$

The shift operator returns 0 if the reference is outside the range of the matrix's index.

**Tensor convolution operator.** This computes a 2D matrix from 3D and 2D matrices. From the pattern matrix W and intensity matrix H, reconstructed matrix U is obtained. The formula is as follows:

$$\mathbf{U} = \mathbf{W} \circledast \mathbf{H} = \sum_{l=0}^{L-1} \mathbf{W}_{\cdot\cdot l} \overset{l\rightarrow}{\mathbf{H}}$$

Each element of U can be calculated as follows, each reconstructed by the sum of k convolutions:

$$u_{nt} = \sum_{k=0}^{K-1} \sum_{l=0}^{L-1} w_{nkl} h_{k(t-l)} \equiv (\mathbf{W} \circledast \mathbf{H})_{nt}$$

**Transpose tensor convolution operator.** This computes a 2D matrix from 3D and 2D matrices. From the pattern matrix W and original matrix V, the overlap matrix R is obtained. The formula is as follows:

$$\mathbf{R} = \mathbf{W} \overset{\top}{\circledast} \mathbf{V} = \sum_{l=0}^{L-1} (\mathbf{W}_{\cdot\cdot l})^{\top} \overset{\leftarrow l}{\mathbf{V}}$$

Each element of **R** can be calculated as follows: each measures the overlap (correlation) between the data and k^th^ neuronal sequence at time t.

$$r_{kt} = \sum_{n=0}^{N-1} \sum_{l=0}^{L-1} w_{nkl} x_{n(t+l)} \equiv \left(\mathbf{W} \overset{\top}{\circledast} \mathbf{V}\right)_{kt}$$

**Math Note S2. Corrections to guarantee convergence of the multiplicative update rules, related to Figure 1.**

**Derivation of the multiplicative update rules.** The error in approximating the original matrix V with the reconstructed matrix U is calculated based on Itakura-Saito divergence as follows:

$$D(\mathbf{V},\mathbf{U}) = \sum_{n=0}^{N-1}\sum_{t=0}^{T-1} \frac{v_{nt}}{u_{nt}} - \log\frac{v_{nt}}{u_{nt}} - 1$$

Because each element of U is calculated from the pattern matrix W and intensity matrix H as follows (see Supplementary Equation 1):

$$u_{nt} = \sum_{k=0}^{K-1}\sum_{l=0}^{L-1} w_{nkl} h_{k(t-l)},$$

differentiating $D(\mathbf{V},\mathbf{U})$ by $w_{nkl}$ or $h_{kt}$ yields the following gradients:

$$\frac{\partial D(\mathbf{V},\mathbf{U})}{\partial w_{nkl}} = \sum_{t=0}^{T-1}\left(\frac{1}{u_{nt}} - \frac{v_{nt}}{u_{nt}^2}\right) h_{k(t-l)}$$

$$\frac{\partial D(\mathbf{V},\mathbf{U})}{\partial h_{kt}} = \frac{\partial}{\partial h_{kt}} \sum_{n=0}^{N-1}\sum_{t=0}^{T-1}\left(\frac{1}{u_{nt}} - \frac{v_{nt}}{u_{nt}^2}\right) = \sum_{n=0}^{N-1}\sum_{l=0}^{L-1} w_{nkl}\left(\frac{1}{u_{n(t+l)}} - \frac{v_{n(t+l)}}{u_{n(t+l)}^2}\right)$$

Furthermore, this can be expressed in matrix form as:

$$\frac{dD(\mathbf{V},\mathbf{U})}{d\mathbf{W}_{\cdot\cdot l}} = \frac{\mathbf{1}}{\mathbf{U}}\left(\overset{l\rightarrow}{\mathbf{H}}\right)^{\top} - \frac{\mathbf{V}}{\mathbf{U}^2}\left(\overset{l\rightarrow}{\mathbf{H}}\right)^{\top}$$

$$\frac{dD(\mathbf{V},\mathbf{U})}{d\mathbf{H}} = \mathbf{W} \overset{\top}{\circledast} \frac{\mathbf{1}}{\mathbf{U}} - \mathbf{W} \overset{\top}{\circledast} \frac{\mathbf{V}}{\mathbf{U}^2}$$

From these gradients and the appropriate learning rates $\eta_{\mathbf{W}}$ and $\eta_{\mathbf{H}}$, the additive update rules are derived.

$$\mathbf{W}_{\cdot\cdot l}^{t+1} = \mathbf{W}_{\cdot\cdot l}^{t} - \eta_{\mathbf{W}} \frac{dD(\mathbf{V},\mathbf{U}^t)}{d\mathbf{W}_{\cdot\cdot l}^{t}} = \mathbf{W}_{\cdot\cdot l}^{t} - \eta_{\mathbf{W}}\left\{\frac{\mathbf{1}}{\mathbf{U}^t}\left(\overset{l\rightarrow}{\mathbf{H}^t}\right)^{\top} - \frac{\mathbf{V}}{(\mathbf{U}^t)^2}\left(\overset{l\rightarrow}{\mathbf{H}^t}\right)^{\top}\right\}$$

$$\mathbf{H}^{t+1} = \mathbf{H}^{t} - \eta_{\mathbf{H}} \frac{dD(\mathbf{V},\mathbf{U}^t)}{d\mathbf{H}^t} = \mathbf{H}^{t} - \eta_{\mathbf{H}}\left(\mathbf{W}^t \overset{\top}{\circledast} \frac{\mathbf{1}}{\mathbf{U}^t} - \mathbf{W}^t \overset{\top}{\circledast} \frac{\mathbf{V}}{(\mathbf{U}^t)^2}\right)$$

Here, by setting $\eta_{\mathbf{W}} = \frac{\mathbf{W}_{\cdot\cdot l}^{t}}{\frac{\mathbf{1}}{\mathbf{U}^t}\left(\overset{l\rightarrow}{\mathbf{H}^t}\right)^{\top}}$ and $\eta_{\mathbf{H}} = \frac{\mathbf{H}^t}{\mathbf{W}^t \overset{\top}{\circledast} \frac{\mathbf{1}}{\mathbf{U}^t}}$, the multiplicative rules are derived.

$$\mathbf{W}_{\cdot\cdot l}^{t+1} = \mathbf{W}_{\cdot\cdot l}^{t} \times \frac{\frac{\mathbf{V}}{(\mathbf{U}^t)^2}\left(\overset{l\rightarrow}{\mathbf{H}^t}\right)^{\top}}{\frac{\mathbf{1}}{\mathbf{U}^t}\left(\overset{l\rightarrow}{\mathbf{H}^t}\right)^{\top}}$$

$$\mathbf{H}^{t+1} = \mathbf{H}^{t} \times \frac{\mathbf{W}^t \overset{\top}{\circledast} \frac{\mathbf{V}}{(\mathbf{U}^t)^2}}{\mathbf{W}^t \overset{\top}{\circledast} \frac{\mathbf{1}}{\mathbf{U}^t}}$$

The multiplicative update rules ensure that the elements of matrices W and H are non-negative. However, the learning rates are fixed in this process, and a case that is too large can be expected, resulting in an increase in the reconstruction error $D(\mathbf{V},\mathbf{U})$ (i.e., $D(\mathbf{V},\mathbf{U}^{t+1}) > D(\mathbf{V},\mathbf{U}^{t})$).

**Reducing the learning rates.** Consider multiplying learning rates by $\alpha$ $(0 < \alpha < 1)$, respectively. The additive update rule for the intensity matrix H is then as follows:

$$\mathbf{H}^{t+1} = \mathbf{H}^{t} - \alpha\eta_{\mathbf{H}}\left(\mathbf{W}^t \overset{\top}{\circledast} \frac{\mathbf{1}}{\mathbf{U}^t} - \mathbf{W}^t \overset{\top}{\circledast} \frac{\mathbf{V}}{(\mathbf{U}^t)^2}\right)$$

$$= (1-\alpha)\mathbf{H}^t + \alpha \times \mathbf{H}^t \times \frac{\mathbf{W}^t \overset{\top}{\circledast} \frac{\mathbf{V}}{(\mathbf{U}^t)^2}}{\mathbf{W}^t \overset{\top}{\circledast} \frac{\mathbf{1}}{\mathbf{U}^t}},$$

where the second term is equal to $H^{t+1}$ when the learning rate is $\eta_{\mathbf{H}}$ (i.e., when $\alpha = 1$), except that $\alpha$ is multiplied. Therefore, since $0 < \alpha < 1$, the second term is non-negative. The first term is also non-negative, and thus the whole remains non-negative. The same holds for the update rule for pattern matrix W. From the above, the following replacements can be applied to reduce the learning rates while guaranteeing that the matrices are non-negative.

$$\mathbf{W}^{t+1} \leftarrow (1-\alpha)\mathbf{W}^t + \alpha\mathbf{W}^{t+1}$$

$$\mathbf{H}^{t+1} \leftarrow (1-\alpha)\mathbf{H}^t + \alpha\mathbf{H}^{t+1}$$

This replacement can be repeated m times to increase the learning rates by a factor of $\alpha^m$, i.e.,

$$(1-\alpha^m)\mathbf{H}^t + \alpha^m\mathbf{H}^{t+1} = (1-\alpha)\mathbf{H}^t + \alpha\{(1-\alpha^{m-1})\mathbf{H}^t + \alpha^{m-1}\mathbf{H}^{t+1}\}$$

We therefore repeated this replacement until $D(\mathbf{V}, \mathbf{U}^{t+1})$ was smaller than $D(\mathbf{V}, \mathbf{U}^t)$.
